# Hck signaling drives long-distance ECM degradation through endo-exocytosis coupling in macrophages

**DOI:** 10.64898/2026.09.03.749091

**Authors:** Cristina Torres-Torres, Marco Ruscone, Satish Babu Moparthi, Lucia Campos-Perello, Anne-Pascale Bouin, Fabrice Senger, Ingrid Bourrin-Reynard, Christiane Oddou, Victoria Pakulska, Yohann Coute, Alexei Grichine, Laurence Calzone, Stéphane Vassilopoulos, Emmanuelle Planus, Olivier Destaing

## Abstract

The diversity of strategies implicated in extracellular matrix (ECM) degradation supporting cell invasion has been poorly achieved. Unlike invasive breast cancer cells, which predominantly degrade the ECM locally via an invadosome-associated degradation, dynamic quantification of live imaging of degradation of physiological ECM such as fibrinogen showed that macrophages employ different modes of ECM degradation since digesting ECM both locally and at long distance. Long distance degradation occurring in macrophage is dependent on both matrix metalloproteases (MMPs) and cathepsins release. Optogenetic manipulation showed that HCK signaling specifically regulates positively this new mode of fibrinogen degradation by increasing cathepsins release and activating localized fusion of acidic CD63-endolysosome vesicles at the vicinity of clathrin hotspots at the rear of macrophages, appearing as a new exo-endocytosis coupling. Computational biology and *in silico* simulations support the importance of the fine spatiotemporal coordination between MMPs and cathepsin activities, and regulation of cathepsin activity by local acid release. Different mode of ECM degradation can thus coexist and this highlight the importance of understanding the cathepsins-MMPs synergy in shaping invasive behavior across different physio-pathological invasion processes.

**“One-sentence summary”:** Macrophage invasion is supported by two distinct modes of ECM degradation, both are partially regulated by HCK signaling, which regulates cathepsins activity by stabilizing acidic vesicles fusion at clathrin hotspots located at the macrophage cell rear.

## Introduction

Cell invasion requires controlled perturbations of highly diverse extracellular environments while preserving the multicellular organization essential for tissue cohesion. In addition to transient interactions with surrounding cells, invasive cells need to migrate through the extracellular matrix (ECM), a non-cellular scaffold that provides structural support and biochemical cues. ECMs can be extremely different in terms of composition, rigidity, porosity and thickness, thereby exposing invasive cells to a wide range of biomechanical challenges. Despite this diversity, the molecular processes implicated in cell invasion have been quite focus on some specialized subcellular structures.

Invadosome superfamily is the ensemble of specialized structures (podosomes, invadopodia, linear invadosomes, etc.) that couple adhesion to ECM degradation, thereby enabling cells to invade surrounding tissues. They share core molecular components and regulatory pathways, and their formation and function strictly depend on actin polymerization, the adaptor Tks5, the small GTPase CDC42 and SRC family kinases activity^1–3^. Invadosomes exert specific 3D mechanical forces on the ECM, which are spatially associated with local ECM degradation supported by proteolytic activity restricted to the immediate vicinity of the invadosomes. The coupling of these two apparently antagonistic functions -adhesion and degradation- is tightly regulated through a mechanism named acto-adhesion and degradation coupling (ADC, ^4,5^). This local degradation is controlled by the local delivery and activation of the transmembrane metalloprotease MT1- MMP (or MMP14) at the vicinity of invadosome, allowing to activate adjacent secreted MMPs^6^. Coupling between ECM degradation and ECM-adhesion structures has also been reported in focal adhesions of some cell typels, which can also display limited ECM-degrading activity under specific conditions^7^.

Invadosomes have been tightly associated with SRC signaling. It is the prototype of tyrosine kinase of the SRC family kinases (SFKs), which comprise eight members. SFKs integrate adhesion, cytoskeletal dynamics, vesicle trafficking, and protease delivery during ECM degradation^8^. Loss-of-function studies revealed an essential role for SRC in invadosome functions in the myeloid lineage, especially osteoclasts and macrophages^9,10^. Moreover, genetic studies showed that HCK kinase was the most redundant kinase to SRC in myeloid cells^11^. Despite being less important for invadosome formation, HCK can still regulates invadosome size and ECM proteolysis in osteoclasts^12^. These findings suggested that SRC and HCK cooperate in matrix degradation but may also possess non-redundant roles. The use of optogenetic version of SRC and HCK (OptoSRC-OS and OptoHCK-OH, respectively) revealed that SRC specifically regulates invadosome dynamics while HCK activates macrophage-specific clathrin endocytosis hotspots^13^.

In addition to the MMP-dependent functions of invadosomes, other extracellular proteases also contribute to ECM degradation, notably the cysteine cathepsins. These pH-dependent proteases are classically confined to the lysosomal/endocytic pathway, where they mediate bulk protein turnover, antigen processing, and pro-protein activation in acidic compartments^14^. However, several cathepsins, particularly cathepsins B, L, S, and K, can be secreted into the extracellular space, where they function as potent ECM-degrading enzymes^14,15^. Once secreted, cathepsins remain inactive as proenzymes (procathepsins) until locally activated under acidic conditions, which are typical of tumors, inflamed tissues, and remodeling sites. Extracellular cathepsins degrade a broad spectrum of ECM components, including collagens, laminins, elastin, and proteoglycans, and can also activate MMPs, amplifying tissue remodeling cascades. Their activity is finely tuned by the local pH and endogenous inhibitors such as cystatins, preventing uncontrolled proteolysis^15^. Among them, Cathepsin B has been shown to localize at invadosomes, where its secretion enhances ECM degradation^16,17^. Yet, despite these observations, the spatiotemporal coordination of extracellular cathepsins with invadosome activity remains poorly understood.

Considering different proteases beyond those classically associated with invadosomes is particularly important because most studies of ECM degradation have relied on fluorescently labeled highly fibrillar collagen I or its hydrolysate, gelatin, as substrates^18,19^. While these systems have been instrumental in defining invadosome function, they overlook how cells remodel non-collagenous matrices, which represent a major fraction of tissue-specific ECM environments. Among these, fibrinogen is a blood-borne glycoprotein cleaved by thrombin to form fibrin, the main structural component of blood clots and provisional matrices at sites of injury^20^. Beyond hemostasis, fibrin and its cleavage products regulate adhesion, migration, proliferation, and inflammation by serving as adhesive scaffolds, reservoirs for growth factors, and substrates for leukocyte trafficking^20,21^. Fibrinogen deposition increases during inflammation, providing a transient matrix that guides cell migration and tissue repair, but also accumulates pathologically in chronic wounds and tumors. In the tumor microenvironment, fibrinogen is abundant, found in 20- 90% of tumor areas^22^, and its degradation product, fibrin, is commonly found at invasive fronts, where it facilitates macrophage infiltration and tumor cell invasion^23^. Macrophages are key effectors of fibrinogen turnover ^24,25^, while also internalizing fibrinogen through Mac-1 (CD11b/CD18) for cathepsin-mediated degradation within lysosomes^26^. Through these complementary mechanisms, macrophages act as central remodelers of fibrin-rich matrices during tissue repair, inflammation, and tumor progression. However, the spatiotemporal and molecular regulation of macrophage-driven fibrinogen degradation remains also poorly understood

To address this question, this study explored macrophage ECM degradation dynamics on fibrinogen, which closely mimics provisional matrices encountered during inflammation and tissue repair. Using long-term live imaging combined with precise spatiotemporal quantification, macrophages were observed to exhibit three distinct modes of ECM degradation: (i) degradation associated with the cell body, (ii) degradation occurring immediately after cell passage, and (iii) a long-distance degradation mode spatially uncoupled from the cell body and correlated with macrophage migration. The coexistence of these distinct modes was further supported by their absence in MDA-MB-231 breast cancer cells, a pathological model of invasive migration that relies predominantly on MMP-mediated proteolysis. Optogenetic activation of OH specifically enhances this process, confirming its functional specificity and causal link to HCK signaling. The long-distance ECM degradation mode dependent on OH is correlated with cathepsins release in the extracellular environment. Mechanistically, HCK promotes this cathepsin-dependent mode of ECM degradation by locally controlling behaviors of acidic vesicles since inducing a novel exo-endocytosis coupling that coordinates the docking and fusion of CD63-positive secretory late endosomes with clathrin hotspots located at the macrophage rear. Exocytosis of these vesicles can mediate local decrease of pH, locally acidifying and activating cathepsin-dependent proteolyzing of the surrounding ECM. In contrast, optogenetic activation of OS fails to trigger this long-distance degradation, underscoring the unique role of HCK in controlling the formation of this specialized exocytic-endocytic domain. Definition of these mode of degradation led us to use computing modeling in order to investigate *in silico* the functional consequences of their interplay. Besides showing the existence of the first exo-endocytosis coupling in macrophages, our approach revealed that different modes of degradation can coexist and opens our understanding of the possible synergic functions of cathepsins and MMPs in numerous physio-pathological conditions.

## Results

### Spatialization and dynamics of ECM degradation activities of Macrophage

The quantitative analysis of ECM degradation dynamics *in vitro*, including changes in degraded area, fluorescence intensity, and degradation rate, remains technically challenging due to obtention of homogenously fluorescently labelled ECM and to perform stable long-term live-degradation imaging (12-20h). In addition, most conventional degradation assays rely on collagen I or its hydrolysate (gelatin), which primarily report MMP-dependent proteolysis and therefore provide limited information on the contribution of other proteolytic systems. To overcome these limitations, a more physiologically relevant matrix composed of fibrinogen mixed with fluorescent-fibrinogen was used to follow ECM degradation by Raw 264.7 macrophages through long-term live-cell imaging (Fig.1A). On this surface, macrophages migrate and actively degrade fibrinogen, revealed by degraded areas appearing in black (Fig.1A,B). To quantify this process in terms of both digested area and intensity of degradation, a computational pipeline was developed to analyze quantitatively the spatiotemporal relationship between ECM degradation and cell migration (Fig.S1A). Precise segmentation of the outside of the cell boundary with the degraded ECM area revealed that degradation occurs not only beneath the cell body but also at sites distant from it. On fibrinogen substrates, macrophages exhibited localized degradation below cell surface (“with cell”, in dark blue), as well as persistent degradation occurring after the cell had migrated away from the digested area (“after cell”, light blue), indicating that fibrinogen proteolysis activity persist at sites previously occupied by the cell (Fig.1C). Moreover, this dynamics analysis showed that ECM degradation events could also occur in regions were the cell never reached during migration (“without cell”, yellow). This observation was further confirmed by kymograph analysis following both the cell border (white line) and digested area (rainbow colors; Fig.1C). This is the first report of long-distance ECM degradation activity by invasive cells. To confirm whether these patterns were specific to fibrinogen, the same analysis was performed on Raw wild-type (wt) macrophages spread on fluorescent gelatin. Compared with fibrinogen, macrophages on gelatin substrates displayed a significantly reduced degradation rate (µm²/min; Fig.1D), which was strictly associated with the cell surface (“with cell”) (Fig.1C,E), being highly localized under the cell. Quantitative analysis revealed that approximately 10% of degradation on fibrinogen occurred at long distance from the cell body, whereas degradation on gelatin was only localized below cell surface (Fig.1E). In this line, the ECM degradation occurring after cell passage (“after cell”) is not directly correlated with the presence of the cell body but represent almost 45% of the fibrinogen degradation activity of macrophage. The separation of degradation fractions and rates for all analyzed cells is shown in Fig.S1B,C. Notably, long-distance degradation on fibrinogen was less intense compared to localized degradation, which was more pronounced, similar to the pattern observed on gelatin (Fig.S1D). Cell position was initially determined from transmitted-light images. Despite phase contrast is highly sensitive to detect membranes and that our computational pipeline designed to conservatively enlarge the segmented cell boundary to account for potential small filopodial extensions potentially missed during segmentation, this same pipe-line of analysis was applied on RAW macrophages expressing a membrane-targeted CAAX fluorescent reporter (RAW-CAAX) in order to observe eventual thin membrane protrusions and filopodia that phase contrast imaging could have missed (Fig.1F). Firstly, comparison of cell segmentation obtained from CAAX fluorescence with the ones obtained from transmitted-light images were not significant for the evaluation of long-distance degradation (Fig.S1E), confirming the occurrence of ECM degradation away from detectable cell-matrix contact. Thus, our computational pipeline clearly confirmed that potential unresolved membrane protrusions are not an explanation for long-distance ECM degradation. Since fibrinogen degradation appears clearly delayed to cell migration, the association between invadosomes and degradation was then investigated (Fig.1G). For that, macrophages expressing Lifeact-RFP were produced and invadosomes were then segmented to be correlated with ECM degradation. On fibrinogen, degradation did not occur massively at invadosome sites but rather immediately after cell passage (Fig.1I), forming a gradient along the cell body that persisted even after the cell had moved away (Fig.1H, S1F,G). In contrast, degradation on gelatin remained consistently associated with invadosomes (Fig.1H, S1F). Importantly, long-distance degradation by macrophages on fibrinogen was not linked to invadosome, since being mostly observed at the rear of the cells (Fig.S1H). Together, quantitative live-cell imaging and these comparative analysis identified three modes of fibrinogen degradation (with, after and without cell) in macrophages characterized by their spatial and temporal correlation, with cell migration. These findings provide evidence that macrophages can employ multiple, spatially distinct modes of ECM degradation.

**Figure 1.**
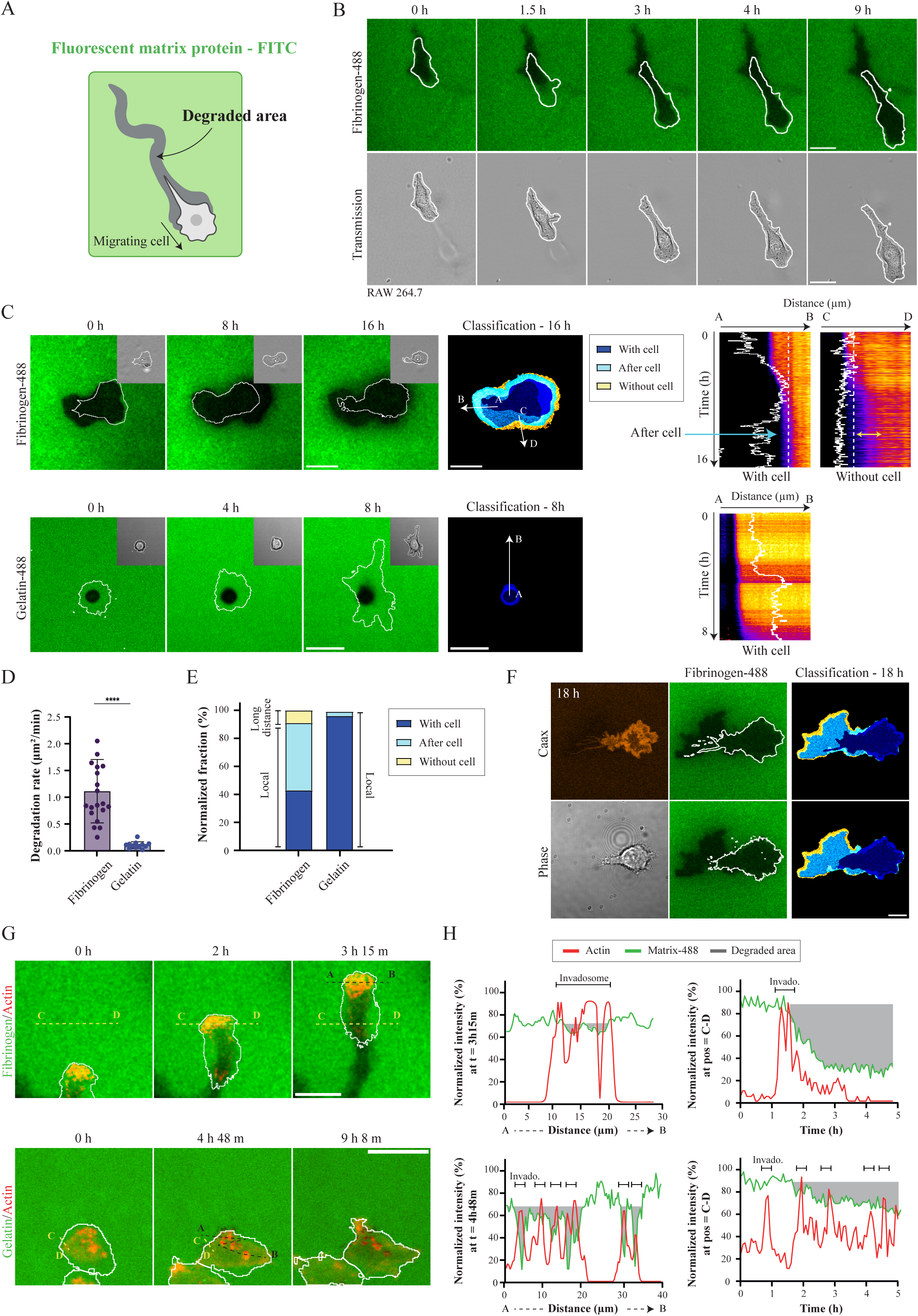
Dynamic degradation of fibrinogen and gelatin by macrophages. (A) Schematic representation of a migrating cell over a fluorescently labeled matrix protein. Degraded areas are shown in black. (B) Representative confocal time-lapse imaging of a Raw 264.7 macrophage migrating on a 2D fluorescent fibrinogen coating, showing localized degradation associated with cell movement. The cell overlay is shown in white. (C) Representative confocal time-lapse imaging of a Raw 264.7 macrophage migrating on 2D fluorescent-fibrinogen or -gelatin coatings (cell overlay white line). Quantitative image analysis showed the classification images indicate 3 degradation patterns associated with the cell (dark blue), degradation occurring after cell passage (light blue), and degradation without cell presence (yellow). Kymographs of matrix degradation (color spectrum) vs the cell border (white line) show its progression in these three categories. Kymograph shows that fibrinogen degradation intensity decreases even at long distance from the cell border (yellow double arrow). (D) Degradation rates (µm²/min) of digesting macrophages on fibrinogen versus gelatin. Data are presented as mean ± SD. Between 20-35 cells per condition were analyzed (N>3). Statistical significance was assessed using an unpaired t test. (E) Quantification of the normalized fraction (contingency plot) of localized versus long-distance degradation on fibrinogen and gelatin. Between 20-35 cells per condition were analyzed (N>3). (F) Representative confocal imaging of a Raw 264.7 macrophage stably expressing the plasma macrophage marker Caax-iRFP migrating on 2D fluorescent-fibrinogen for 18h. Analysis of classification images still indicate the same 3 degradation patterns using another way to follow cell border. (G) Representative confocal time-lapse imaging of invadosome (lifeact-RFP) association with degradation on fibrinogen and gelatin. (H) At a given time point, colocalization of invadosome with the matrix is shown (normalized mean intensity as a function of distance). Temporal progression of degradation at the same sites is shown (normalized mean intensity as a function of time). For all time-lapse sequences, frames were acquired every 4 minutes for up to 16 hours. Scale bar: 20 µm.

### Delayed and long-distance degradation are specific features of macrophages

To determine whether the different modes of ECM degradation were specific to macrophages or represented a more general feature of invasive cells, macrophages were compared with highly invasive MDA-MB-231 breast cancer cells, a widely used model of invadosome activity and cancer cell invasion. Their degradation behavior was then analyzed on both fibrinogen and gelatin substrates. On fibrinogen, MDA-MB-231 cells exhibited degradation forming an intracellular gradient, extending over time from the cell periphery toward the cell center, as observed in both fluorescence images and kymographs (Fig.2A). In contrast, degradation on gelatin appeared as punctate areas also beneath the cell body. No significant differences in degradation rates (µm²/min) were observed between MDA-MB- 231 cells and Raw macrophages on either fibrinogen or gelatin (Fig.2B), indicating that the rate of matrix digestion was comparable between both cell types. However, for both cell type, the degradation rate was markedly lower on gelatin than on fibrinogen, reflecting a more spatially restricted degradation pattern. On fibrinogen, degradation instead extended over a larger fraction of the cell-associated area. All quantitative analyses were performed exclusively on cells that actively degraded the substrate, excluding non-degrading cells from the dataset. Qualitative classification further revealed that MDA-MB-231 cells exhibited local degradation limited to their cell body on both fibrinogen and gelatin substrates, unlike macrophages on fibrinogen, which uniquely displayed delayed and long-distance degradation modes on fibrinogen (Fig.2C). To determine the relationship between degradation and invadosomes, MDA-MB-231 cells expressing LifeAct-RFP were analyzed on both matrices. On fibrinogen, degradation was initially spatially associated with invadosomes, starting at the leading edge and progressively extending inward in a gradient-like manner (Fig.2D,E, S2A). In contrast, degradation on gelatin remained strictly associated with single invadosomes (Fig.2D,E, S2A).

**Figure 2.**
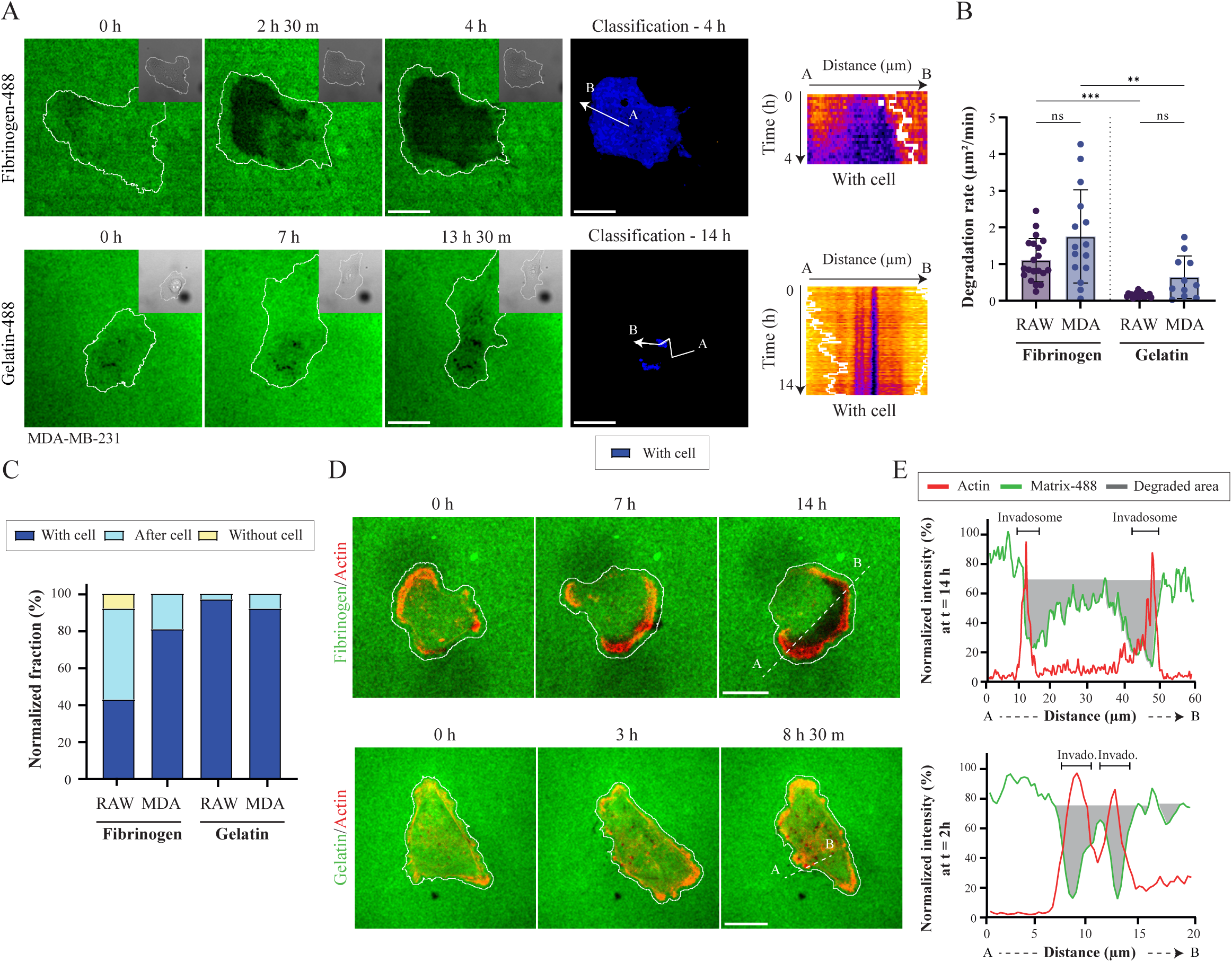
Comparative degradation dynamics of MDA cancer cells and macrophages on fibrinogen and gelatin substrates. (A) Representative confocal time-lapse imaging of MDA cells on fibrinogen, showing intracellular gradient-like degradation, versus punctate degradation on gelatin. Classification images indicate degradation associated with the cell (dark blue). Kymographs of matrix degradation depict its progression. The white line marks the cell border. (B) Degradation rates (µm²/min) of MDA cells on fibrinogen versus gelatin, compared with Raw cells. Data are presented as mean ± SD. Between 10-35 cells per condition were analyzed (N>3). Statistical significance was assessed using one-way ANOVA. (C) Quantification of the normalized fraction (contingency plot) of localized versus long-distance degradation in MDA cells compared with Raw. Between 10-35 cells per condition were analyzed (N>3). (D) Representative confocal time-lapse imaging of RAW 264.7 macrophage stably expressing LifeAct-RFP to follow invadosome-associated degradation on fibrinogen versus on gelatin. (E) At a given time point, colocalization of lamellipodia or invadosome with the matrix is shown (normalized mean intensity as a function of distance). For all time-lapse sequences, frames were acquired every 10 minutes for up to 16-24 hours. Scale bar: 20 µm.

The absence of long-distance degradation-after cell and without cell- in MDA-MB-231 cells, together with the close temporal association between invadosomes and ECM degradation on both fibrinogen and gelatin, supports the existence of distinct degradation modes. The long-distance one appears specific to macrophages.

### SRC family kinases regulates differently these mode of fibrinogen degradation and their dynamics in macrophage

The identification of these distinct ECM degradation modes led to investigate their regulatory mechanism. Taking advantage of previously developed optogenetic tools that allow positive or negative activation of macrophage migration^13^, the effects of optoSRC (OS) and optoHCK (OH) activation were determined in RAW macrophage stably expressing these constructs. Upon activation, OS macrophages displayed local degradation patterns comparable to those observed in wt macrophages (exposed to the same blue light illumination protocol; Fig.3A-C). In contrast, OH activation markedly shifted the degradation fraction toward the long-distance mode, accompanied by a reduction in “with cell” mode (Fig. 3A-C). Quantitative classification revealed that activated OH macrophages degraded more than 30% through a long-distance mode, which is associated with a significant increase in the degradation rate of this specific mode (Fig.3B; in yellow) compared to OS or wt macrophages. The distribution of degradation fractions and degradation rates per cell, stratified by degradation mode and condition (wt, OS, OH), are shown in Fig.S3A-B. Notably, the intensity of long-distance degradation was markedly increased in OH macrophages, whereas the intensity of local degradation remained similar between OS and OH (Fig.S3C).

**Figure 3.**
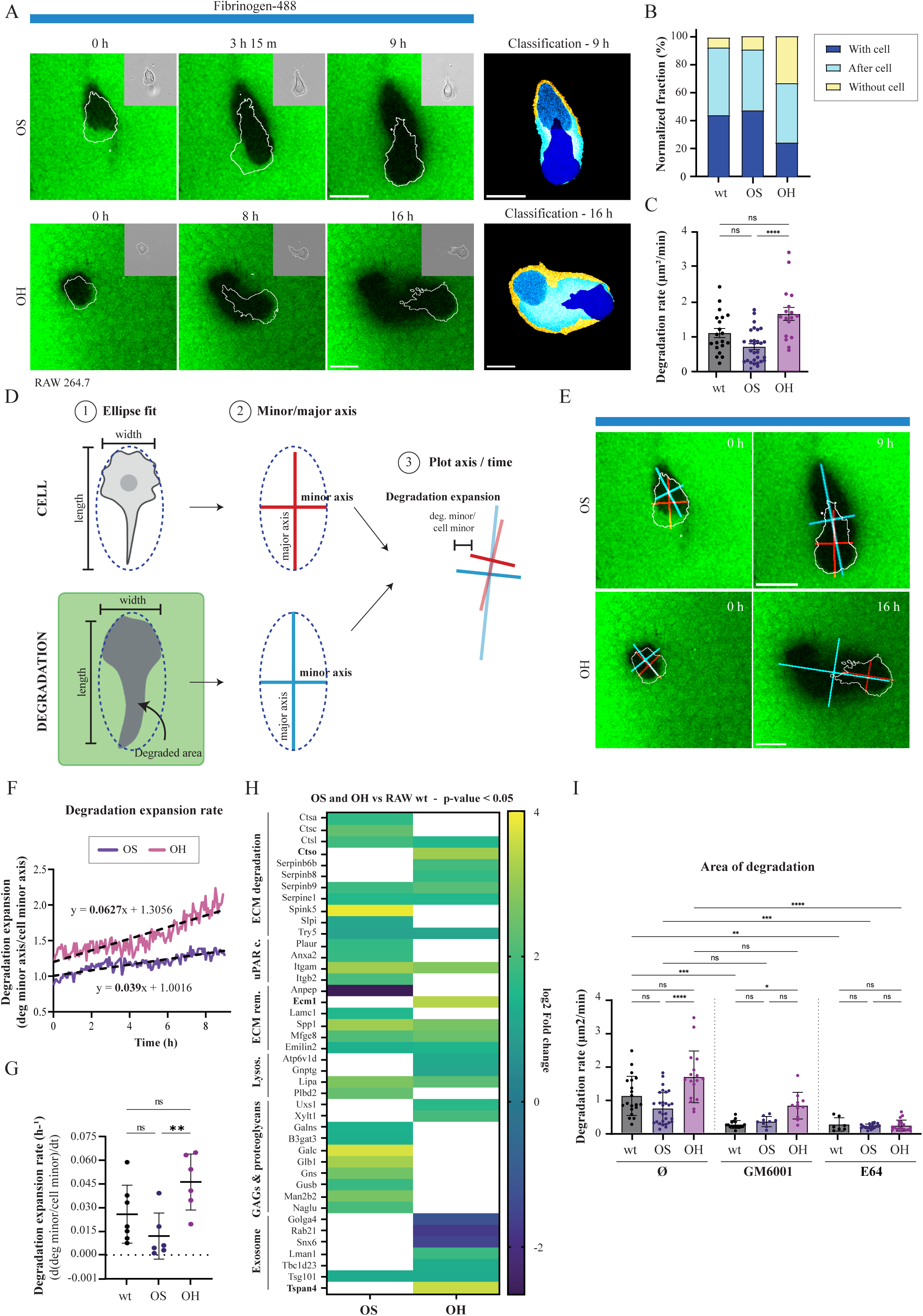
Role of Src family kinases in macrophage degradation dynamics. (A) Representative confocal time-lapse imaging of Raw 264.7 macrophages expressing OS or OH on fibrinogen coatings and activated by blue light imaging (4,16 mHz, 1 image every 240s). Classification images indicate degradation associated with the cell (dark blue), after cell passage (light blue), or without cell presence (yellow). (B) Quantification of the normalized fraction (contingency plot). Between 20-30 cells per condition were analyzed (N>3). (C) Degradation rates (µm²/min) of RAW macrophages (wt, OS or OH) on fibrinogen and photoactivated with blue light (4,16 mHz). Data are presented as mean ± SD. Between 20- 30 cells per condition were analyzed (N>3). Statistical significance was assessed using one- way ANOVA. (D) Schematic representation of the degradation expansion analysis. Cell borders and ECM degradation areas were independently fitted with ellipses defined by major and minor axes. Differences between the dimensions of the cell and degraded area were used to assess the spatial uncoupling between cell migration and ECM degradation. To quantify lateral degradation expansion independently of cell width, the minor axis of the degraded area was normalized to the minor axis of the corresponding cell (degradation minor axis/cell minor axis). (E) Representative confocal time-lapse images of RAW 264.7 macrophages expressing OS or OH migrating on fibrinogen and photoactivated with blue light. Ellipse fitting was used to follow the spatial evolution of the cell and degraded area over time. While extension along the major axis largely reflected the direction of cell migration, expansion along the minor axis revealed degradation extending laterally beyond the cell boundaries, particularly following OH activation (F) Representative time course of degradation expansion in Raw macrophages expressing OS or OH. Degradation expansion was calculated as the ratio between the minor axis of the degraded area and the minor axis of the corresponding cell. Linear regression was used to determine the degradation expansion rate. (G) Quantification of expansion rates in wt, OS-, and OH-expressing macrophages (h□¹). Data are presented as mean ± SD. Between 6-7 cells were analyzed per condition from more N>3. Statistical significance was assessed by one-way ANOVA. (H) Secretome heatmap showing significantly regulated proteins (p ≤ 0.05) plotted as log□fold change (log□FC). Data were obtained from triplicates after 24 h stimulation in serum- free conditions. (I) Degradation rates (µm²/min) of RAW macrophages (wt, OS or OH) on fibrinogen treated with a pan-MMPs inhibitor (GM6001, 10 µM) or a pan-cathepsin inhibitor (E64, 50 µM). Data are presented as mean ± SD. Statistical significance was assessed using a Kruskal-Wallis test. Between 10-30 cells per condition were analyzed. For all time-lapse sequences, frames were acquired every 4 minutes (photoactivation is intrinsically done 4.16 mHz) for up to 16 hours. Scale bar: 20 µm.

To further demonstrate that long-distance degradation reflects the progressive spreading of proteolytic activity beyond the cell, a geometrical analysis was developed to quantify the expansion of the degraded area over time. In real time, ellipses were independently fitted to the segmented cell body area and the associated degraded region. Ellipses were then defined by their major and minor axes of both structures evolving over time (Fig.3D). Analysis of each ellipses showed that the degraded region progressively extended beyond the cell body in both axes (Fig. S3D,E). Along the major axis, the degradation area extended beyond the length of the cell body, consistent with the accumulation of degradation along the migration path. Importantly, the minor axis also progressively increased beyond the cell width, demonstrating lateral expansion of degradation that could not be explained solely by cell displacement.

Thus, this geometrical analysis was applied on both macrophages where OS or OH were activated during degradation time (Fig.3E). At the end of the acquisition (9 h), the degraded region extended ∼6 µm beyond the total cell width in OS macrophages and up to ∼20 µm in OH macrophages, mainly due to a symmetrical lateral extension of degraded area (approximately 3 µm for OS- and 9-10 µm for OH-macrophages, respectively). Because changes in cell morphology could influence the absolute dimensions of the degraded area, lateral degradation expansion was normalized to cell size by calculating the ratio between the minor axis of the degraded region and the one of the cell body. The evolution of this ratio over time therefore provided a measure of degradation expansion beyond the cell boundary independently of changes in cell width. This analysis revealed progressive lateral expansion of the degraded area in both OS- and OH-expressing macrophages (Fig.3F, S3D). However, degradation expansion presented significant different kinetics since OH activation promoted a substantially faster increase in the degradation-to-cell minor-axis ratio than during OS activation. Linear regression of these curves was then used to extract a *degradation expansion rate* for each cell (Fig. 3G). Representative regression slopes corresponded to approximately 0.039 h□¹ for OS- and 0.063 h□¹ for OH-macrophages. These results were consistent with the increased long-distance degradation observed upon HCK activation. Together, these measurements provide quantitative evidence that the increase in degraded area cannot be explained solely by changes in cell morphology or migration, but reflects a progressive expansion of fibrinogen degradation beyond the cellular boundaries.

To elucidate the mechanisms underlying OH-dependent long-distance degradation, secretome analysis was performed by comparing the protein secretion profiles of WT macrophages with those of OS- and OH-stimulated macrophages (Fig. 3H). Proteins with a p-value ≤ 0.05 (−log10(p-value) ≥ 1.3) and a log2 fold change ≥ 1 were considered differentially secreted. Shared and condition-specific proteins are shown in Fig. S4A-C. The secretome revealed distinct ECM-remodeling profiles between OS and OH macrophages. Although no MMPs were identified among the differentially secreted proteins, OS macrophages displayed changes in proteins associated with cell adhesion and pericellular ECM remodeling, including Plaur, Anxa2, Itgam and Itgb2, consistent with the predominantly local degradation phenotype observed upon SRC activation. In contrast, OH macrophages showed a stronger representation of proteins associated with lysosomal and vesicular pathways, including cathepsin O, Atp6v1d and Tspan4, together with several serpins. Cathepsin L was detected in both OS and OH conditions. OH macrophages also showed changes in ECM- and adhesion-associated proteins, including Ecm1, while Tsg101 was shared between both conditions (Fig. 3H). Consistently, Gene Ontology enrichment analysis of cellular components revealed enrichment of lysosomal, vacuolar, and vesicle-related proteins in both OS and OH macrophages (Fig. S4C).

The secretome profile therefore suggested that the HCK-dependent long-distance degradation phenotype could rely preferentially on cathepsin- and lysosome-associated proteolytic activity rather than on a classical MMP-dominated mechanism. To functionally test this hypothesis, live-cell fibrinogen degradation assays were performed in the presence of the broad-spectrum MMP inhibitor GM6001 or the cysteine cathepsin inhibitor E64 (Fig.3I). Inhibition of each family of proteases reduced fibrinogen degradation in wt, OS, and OH macrophages, confirming that both MMPs and cathepsins contribute to matrix degradation. However, OH-activated macrophages were markedly more sensitive to cathepsin inhibition than to MMP inhibition, supporting a predominant contribution of cathepsins to the HCK-dependent long-distance degradation mode. Consistently, E64 also impaired macrophage fibrinogen degradation, whereas GM6001 strongly reduced, but did not completely abolish, it.

### OH activation promotes exocytosis of CD63-positive endolysosomal compartments at the macrophage rear

In order to understand how OH activation could promote cathepsin-dependent long-distance degradation at macrophage rear, the dynamics of acidic endolysosomal compartments at the plasma membrane were investigated. Indeed and on the contrary with MMPs, the activation of most cathepsins is dependent on an optimal range of acidic pH (active below 7 but inhibited at pH3)^27^. Previous work showed that activated OH is specifically relocalized with clathrin endocytic hotspots at the rear of migrating macrophages, but not with both Lamp1 or Lamp2 vesicles^13^. Thus, CD63-positive vesicles were monitored by TIRF microscopy, allowing visualization of vesicles at the vicinity of the plasma membrane where activated OH is relocalized in specific subcellular structures^13^. Upon light activation, activated OH is relocalized to an immobile subcellular structure (indicated by strain of activated OH on the kymograph) that appears also accumulating CD63-GFP-positive vesicles also immobile (Fig.4A,D). Kymograph and fluorescence-intensity analyses revealed repeated and transient recruitment of CD63-GFP-positive vesicles to OH-enriched sites, suggesting a dynamic association between HCK signaling domains and endolysosomal vesicles. To determine whether these docked CD63-positive vesicles associated with OH could also be the sites of vesicles fusion with the plasma membrane, CD63-pHluorin was expressed in macrophages stably expressing OH. Indeed, the fluorescence of the pHluorin is quenched by the acidic content of CD63 vesicles and increases drastically when the acidic lumen of CD63-positive vesicles becomes exposed to the neutral extracellular environment. CD63-pHluorin bursts repeatedly occurred in close spatial and temporal association after activation of OH and at the vicinity of OH-positive sites (Fig.4B). Consistently, VAMP8, a SNARE involved in late endosomal/lysosomal membrane fusion, was also transiently recruited to OH-positive hotspots following HCK activation (Fig.4C). To confirm the activation of CD63-vesicles fusion downstream OH activation, co-occurrence analysis was performed to correlate formation and activation of OH with different fusion markers (Fig.4D). The strongest associations of OH spots were observed with VAMP8 (39.3%), CD63-GFP (37.8%), CD63- pHluorin (35.8%), and ATG9-pHluorin (31.2%). These associations were strongly polarized toward the macrophage rear. Moderate associations were also detected with vATPaseOc (24.0%), TSG101 (22.8%), and VAMP7 (20.8%), whereas Rab8b (15.2%) and synaptotagmin V (11.8%) showed lower co-occurrence. Although ATG9 is classically associated with autophagy, no detectable association was observed with the autophagosomal marker LC3 (data not shown), suggesting that the ATG9-positive structures identified here may instead reflect non-canonical endolysosomal trafficking. Thus, activation of HCK signaling induced the docking and the fusion of acidic CD63- positive vesicles that should decrease locally the pH of the extracellular environment, probably mostly at macrophage rear. In addition to modify behaviors of acidic vesiciles, the association of activated OH with cathepsins was also tested. However, following cathepsin dynamics is challenging since probably highly transient. Few available GFP-tagged cathepsins were tested and the fluorescent CtsL-GFP was used primarily to follow their localization in response to OH activation. In wt and activated OS macrophages, CtsL-GFP was mainly detected in transient vesicles and mostly recruited at the middle of the invadosome ring (white arrows, Fig.4E). In OH macrophages, rare association of CtsL-GFP and activated OH spots were observed at the rear of macrophages (white arrow, Fig.4E). These data rather support that activation of long distance degradation of OH is mostly associated by its ability to induce fusion of acidic CD63-positive vesicles with the plasma membrane at the macrophage rear. Thus, the functional requirement for maintaining low pH in vesicular compartments in OH-dependent degradation was assessed. Inhibition of v-ATPase with bafilomycin A1 markedly reduced fibrinogen degradation in OH-activated macrophages, while having a substantially weaker effect in wt and OS cells (Fig.4F,G). These results confirmed that the enhanced degradative activity induced by HCK depends on acid intracellular CD-63 positive vesicular compartments.

**Figure 4.**
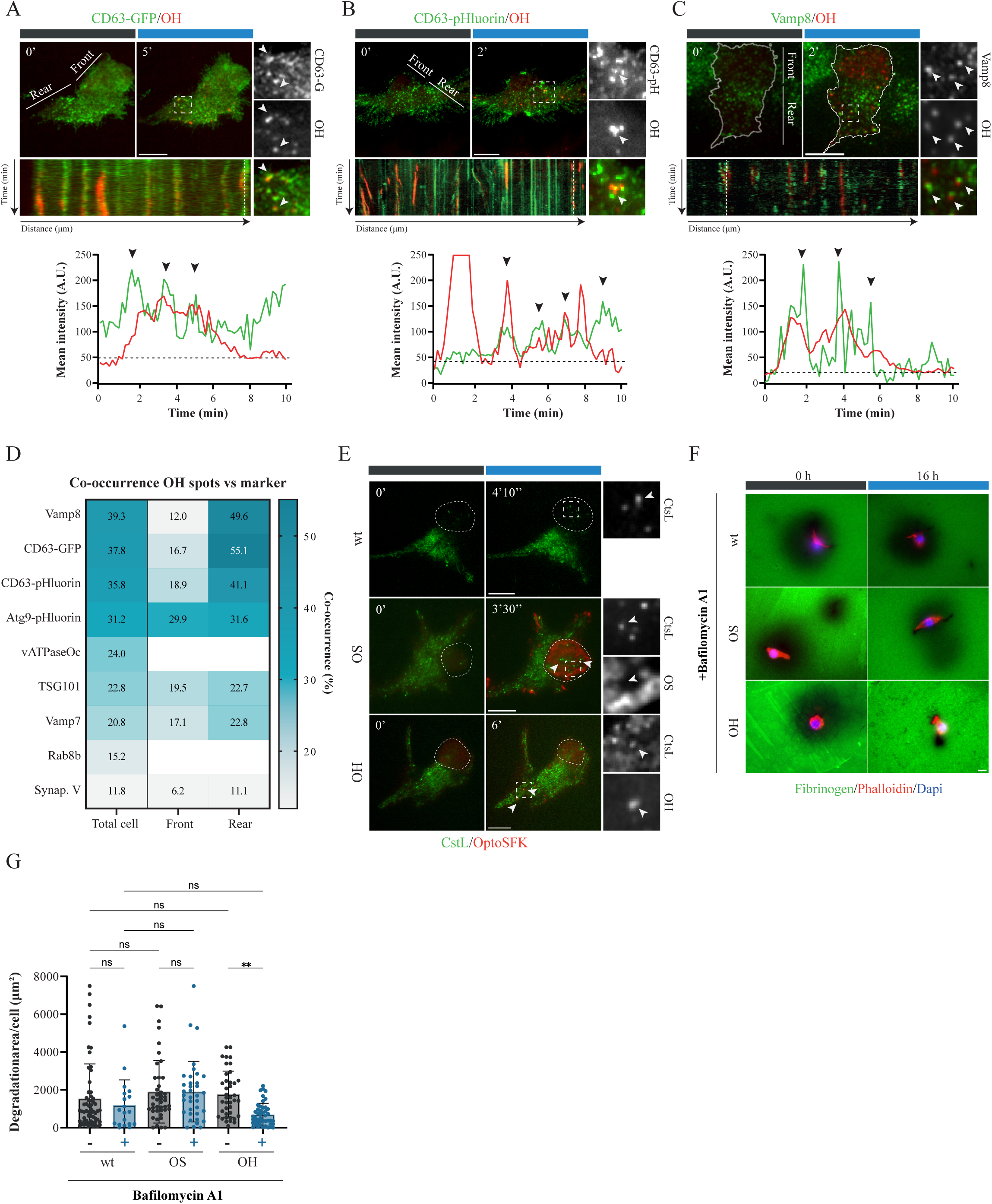
OH-mediated docking and exocytosis of acidic CD63 vesicles at the plasma membrane of macrophages drives long-distance macrophage degradation. (A-C) Representative time-lapse TIRF imaging of CD63-GFP, CD63-pHluorin, and VAMP8-GFP in OH macrophages. Kymographs at the rear show hotspots (repeated CD63 and Vamp8 accumulations) colocalizing with OH. Temporal patterns are shown as mean intensity/time. (D) Co-occurrence analysis of OH spots with numerous vesicle markers. Between 5-17 cells per condition were analyzed. (E) Representative time-lapse TIRF imaging of wt, OS, and OH Raw 264.7 macrophages expressing CtsL-GFP. Frames were acquired every 10 seconds for 10 minutes. Scale bar: 5 µm. (F) Representative immunofluorescence of Raw 264.7 wt, OS, and OH cells light stimulated or not (blue and grey box) treated with bafilomycin A1 (50 nM) for 16h. (G) Degradation areas per cell (µm²) of RAW macrophages (wt, OS or OH) on fibrinogen photoactivated or not (4,16 mHz) and treated with Bafilomicin A1 (50nM). Data are presented as mean ± SD. Between 20-40 fields per condition were analyzed, n > 3. Statistical significance was assessed using one-way ANOVA.

Combined with the dependence of long-distance degradation on cathepsin activity and vesicle acidification, these observations support a model in which HCK promotes spatially controlled release of acidic, protease-containing compartments at the macrophage rear.

### OH activation reveal a unique exocytosis-endocytosis coupled mechanism that sustains controlled membrane fusion of CD63 endolysosome compartment

The close association between activated HCK (OH), clathrin endocytic hotspots, and CD63- positive fusion events raised the possibility that endocytosis and endolysosomal exocytosis could be functionally coupled. Thus, the contribution of acidic vesicle function to this association was first assessed by inhibiting low pH in CD63 vesicles, using bafilomycin A1. v-ATPase inhibition strongly reduced the co-occurrence between OH-positive sites and CD63-pHluorin events and disrupted their coordinated dynamics (Fig.5A,B), indicating that the acidity state of the endolysosomal compartment is required for its association with activated HCK. Our previous work showed that HCK activation promotes actin-dependent clathrin endocytic hotspots at the rear of migrating macrophages^13^. To determine whether this endocytic activity also contributes to the regulation of CD63-positive exocytosis, clathrin-mediated endocytosis was inhibited using jasplakinolide, which stabilizes F-actin and suppresses membrane fission in clathrin hotspots induced by OH. Despite inhibition of productive endocytosis, a strong spatial association between OH and CD63-pHluorin-positive compartments was maintained (Fig.5A,B). However, the characteristic transient CD63-pHluorin bursts observed under control conditions (Fig.4A) were lost, and CD63- positive structures instead accumulated at the plasma membrane in association with OH (plateau, Fig.5A). These observations suggest that OH-dependent endocytic dynamics are not required for the initial recruitment of CD63-positive compartments, but are necessary for their normal turnover at the membrane.

**Figure 5.**
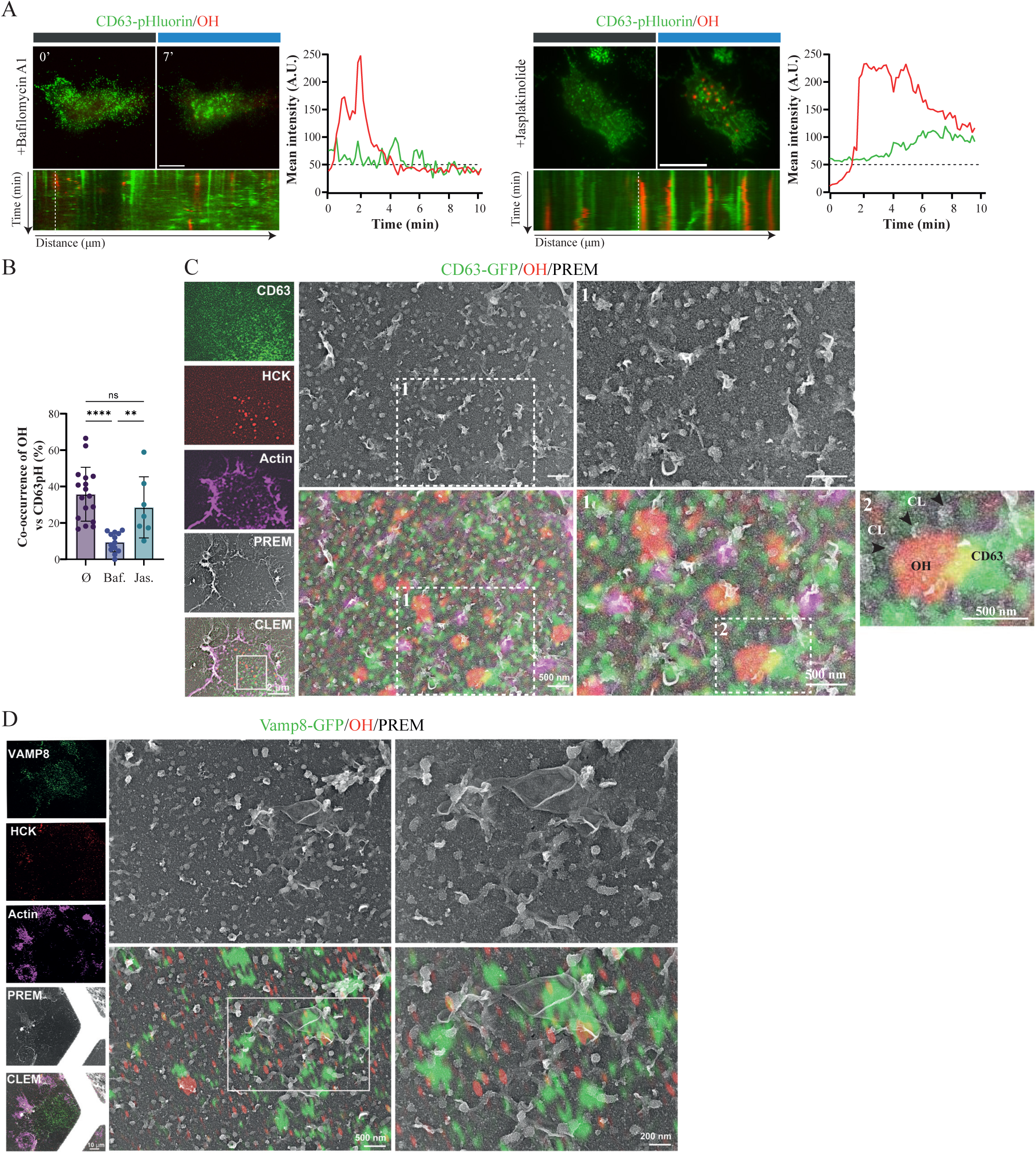
OH-mediated exocytosis-endocytosis coupling between acid CD63 vesicles docked at the plasma membrane and clathrin hotspots. (A) Representative time-lapse TIRF imaging of CD63-pHluorin in OH macrophages treated with either bafilomycin A1 (50 nM) and jasplakinolide (0.5 µM). Loss of acid identity of CD63 vesicles after bafilomycin A1 treatment decreases association and fusion of these vesicles at activated OH spots, whereas the endocytosis fission inhibitor jasplakinolide traps vesicles at the membrane. Kymographs and temporal patterns are shown as mean intensity over time. (B) Co-occurrence of OH spots with CD63-pHluorin under bafilomycin or jasplakinolide treatment. Data are presented as mean ± SD. Between 7-17 cells per condition were analyzed. Statistical significance was assessed using one-way ANOVA. (C-D) CLEM image showing OH (red), clathrin (CL, black arrow), and CD63-GFP or Vamp8 (green) in a coupled exo/endocytosis structure. Scale bar: 500 nm. Frames were acquired every 10 seconds for 10 minutes stimulation. Scale bar: 5 µm.

To further examine the spatial organization of of both CD63-vesicles exocytosis (Fig.4) and OH-dependent clathrin hotspots endocytosis^13^ at the ultrastructural level, correlative light and electron microscopy (CLEM) was performed following plasma membrane unroofing. Activated OH-positive domains were frequently surrounded by clathrin-coated structures, consistent with previously described association of OH with clathrin hotspots^13^. Strikingly, CD63-GFP-positive membrane structures were detected in close proximity to these OH/clathrin domains (Fig.5C), and a similar spatial association was observed for VAMP8- GFP-positive structures, OH and clathrin pits (Fig.5D). These observations support that activation of OH can physically couple both exocytosis of CD63- endolysosomal vesicles and some endocytic clathrin structures, at the nm-scale. Importantly, the CLEM data did not allow direct visualization of vesicle fusion events. Plasma membrane unroofing may disrupt or remove transient membrane intermediates, and the temporal scales of the two processes are markedly different: exocytic fusion occurs rapidly (seconds), whereas clathrin-mediated endocytosis proceeds over substantially longer timescales (minutes). Thus, the ultrastructural data should be interpreted as evidence of spatial proximity rather than direct morphological demonstration of simultaneous exo-endocytic events.

Together, the pharmacological and correlative imaging data support a functional coupling between HCK-dependent clathrin endocytosis and endolysosomal exocytosis at the macrophage rear. HCK appears to organize a membrane-remodeling domain in which CD63- and VAMP8-positive compartments are recruited and undergo controlled fusion in close proximity to clathrin endocytic hotspots. Such coupling could provide a mechanism to locally release acidic vesicular content and cathepsins while simultaneously maintaining plasma membrane homeostasis, thereby sustaining spatially restricted long-distance ECM degradation.

### Computational modeling recapitulates spatially distinct modes of macrophage-mediated ECM degradation

To challenge the functional importance of these different modes of degradation, an agent-based computational model was developed using PhysiCell to simulate a migrating and degradative. The model incorporated experimentally derived secretion rates and decay constants for each component (Material and methods). In addition to test a limited number of core elements, this computational model allows testing numerous possible activation patterns. A migrating macrophage was represented as a polarized cell in which MMP activity was restricted to the leading edge, whereas cathepsins were released all over the cell, and acids (H+) were released extracellulary from the rear (Fig.6A, S5A). As a baseline control, a non-migrating, non-degrading cell produced no detectable ECM degradation over 24 h, confirming the absence of spontaneous matrix loss in the model (Fig. S5C). Cathepsin-mediated ECM degradation was modeled as dependent on local H+ availability, while MMP-mediated degradation occurred locally at the cell front. The corresponding secretion, diffusion, decay, and ECM-interaction parameters were incorporated into the simulation framework (Fig.6B, S5A,B). When MMP activity was simulated alone, ECM degradation remained highly localized and closely followed the migrating cell, reproducing a predominantly cell-associated degradation pattern (Fig.6C). In contrast, cathepsin and H+ release generated a broader degradation field extending behind the migrating cell. Combining both systems reproduced the main spatial features observed experimentally, with localized degradation associated with the cell front together with a broader rear- associated degradation pattern (Fig.6C). These simulations therefore support the concept that the balance between locally acting MMPs and diffusible acid-dependent cathepsin activity can determine the spatial organization of ECM degradation.

**Figure 6.**
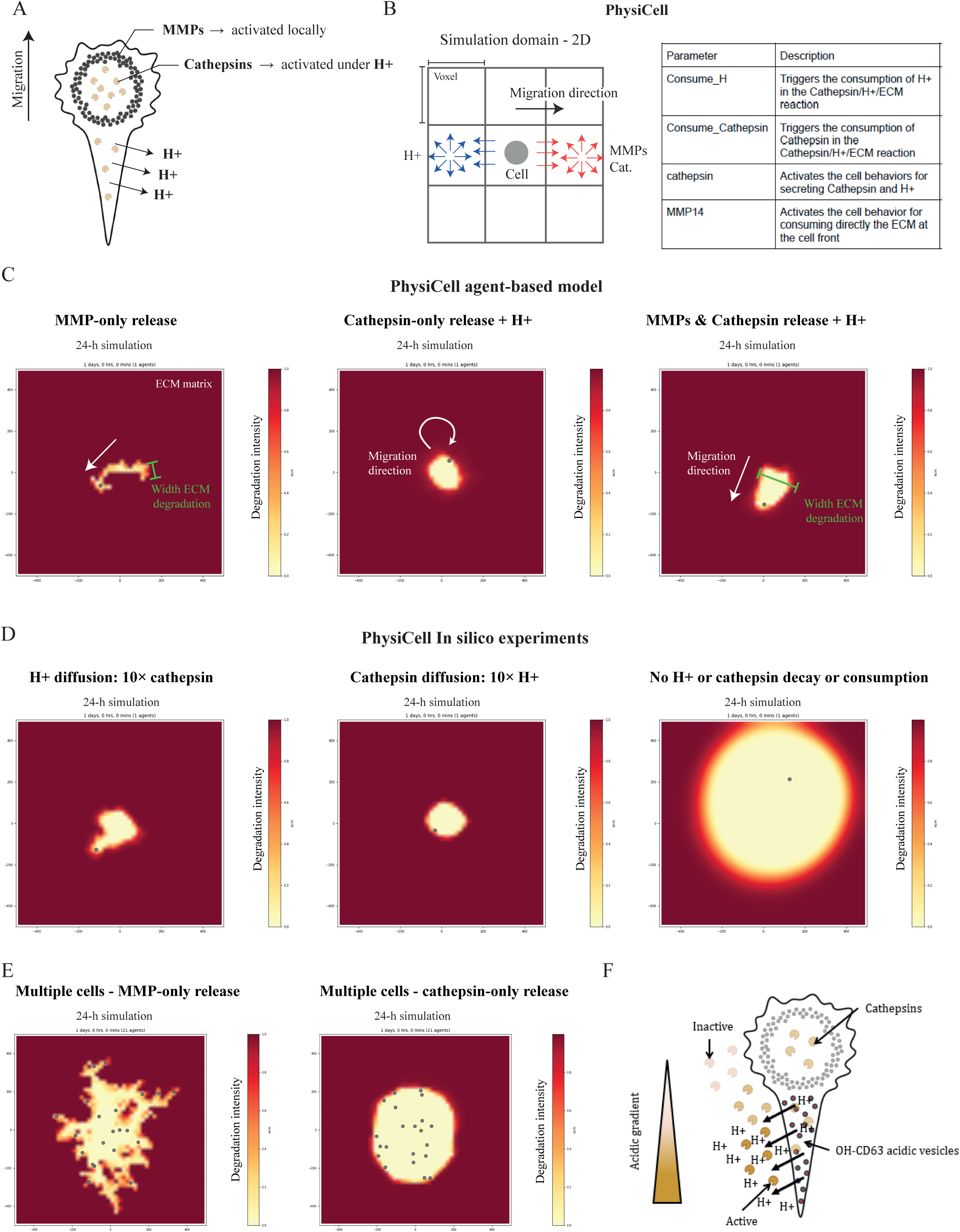
Computational modeling of spatially distinct macrophage-mediated ECM degradation modes. (A) Schematic representation of the polarized proteolytic model used for simulations. MMP activity was restricted to the leading edge of the migrating cell, whereas cathepsins and H+ were released at the cell rear. Cathepsin-mediated ECM degradation was dependent on local H+ availability. (B) PhysiCell agent-based model implemented in a two-dimensional simulation domain. The migrating cell interacts with neighboring voxels containing diffusible H+ and cathepsins, while MMP-mediated ECM degradation is locally restricted to the cell front. Main cellular behaviors implemented in the model are indicated. (C) Representative 24-h simulations of ECM degradation generated by MMP-only release, cathepsin release combined with H+, or simultaneous MMP and cathepsin/H+ activity. MMP activity generated degradation closely associated with the migrating cell, whereas cathepsin/H+ release produced broader degradation extending from the cell rear. Combination of both proteolytic systems reproduced localized and long-distance degradation patterns. Green bars indicate the width of ECM degradation. (D) In silico perturbation of H+ and cathepsin diffusion and turnover. Representative 24-h simulations are shown for H+ diffusion increased 10-fold relative to cathepsin diffusion, cathepsin diffusion increased 10-fold relative to H+ diffusion, and absence of H+ and cathepsin decay or consumption. Increased H+ diffusion broadened the degradation pattern, whereas unrestricted persistence of both components resulted in extensive ECM degradation throughout the simulation domain. (E) Representative 24-h simulations of multicellular ECM degradation. Cells relying exclusively on MMP-mediated degradation generated irregular, cell-associated degradation tracks, whereas cathepsin-dependent degradation produced a broader and more continuous degradation field around the cell population. (F) Proposed model of macrophage long-distance ECM degradation. Rear-localized fusion of OH-associated CD63-positive acidic vesicles releases H+ and cathepsins into the extracellular environment. The resulting acidic gradient progressively decreases with distance from the cell, spatially restricting cathepsin activity and ECM degradation.

The model was next used to explore how the relative diffusion properties of H+ and cathepsins influence the degradation pattern. Increasing H+ diffusion tenfold relative to cathepsins broadened the degradation field, whereas increasing cathepsin diffusion tenfold relative to H+ produced a more spatially restricted pattern (Fig.6D). Importantly, removal of H+ and cathepsin decay or consumption resulted in uncontrolled expansion of ECM degradation throughout the simulation domain, demonstrating that degradation range critically depends on the finite lifetime and local consumption of both components (Fig.6D).

Finally, this basic model was extended to multicellular environments in order to explore how distinct proteolytic programs could shape matrix remodeling at larger spatial scales and in a multicellular environment, as mostly reported for macrophages in invaded tissues. Populations relying exclusively on MMP-mediated degradation generated irregular, cell-associated degradation tracks, whereas cathepsin-dependent degradation produced a broader and more continuous degradation field around the cell population (Fig.6E). These simulations illustrate how the relative contribution of local versus diffusible proteolytic mechanisms could generate markedly different patterns of tissue-scale ECM remodeling. Together, the computational model recapitulates the experimentally observed coexistence of local and long-distance degradation modes and supports a mechanism in which spatially restricted MMP activity is complemented by rear-localized release of H+. In this framework, the diffusion and lifetime of acidic and proteolytic components determine the spatial range of degradation, providing a mechanistic basis for the extended ECM degradation observed in macrophages (Fig.6F).

## Discussion

Quantitative analysis of live ECM degradation by macrophages revealed that different modes of degradation can coexist. Fibrinogen degradation occurs both at the level of the cell body (locally) and at sites distant from it, either after cell passage or at long distances from the cell. Pharmacological approaches showed that the newly identified long-distance mode of fibrinogen degradation depends on cathepsin activity and acidic vesicular compartments and is specifically enhanced by an optogenetic activation (OH) of HCK signaling. This OH-dependent exocytosis (docking and fusion) of CD63-positive endolysosomal provides a potential mechanism for localized cathepsin release and extracellular acidification at the macrophage rear, where long-distance degradation occurs. This process is tightly regulated and functionally linked to clathrin-mediated endocytosis, supporting the existence of an HCK-regulated exo-endocytic coupling mechanism. To our knowledge, this represents a previously undescribed mode of spatial coordination between endocytosis and endolysosomal exocytosis in macrophages. Computational modeling of both ECM proteolytic systems further supported the idea that distinct degradation modes generate different spatial patterns of matrix remodeling at both the single-cell and multicellular levels.

### Multiple spatialization of ECM degradation

Precise quantification of live-cell imaging data from macrophages migrating on degradable substrates revealed distinct ECM degradation behaviors. Besides their specific rates, one characteristic of these different behaviors is their distinct spatial distribution. Long-distance degradation and degradation persisting after cell passage demonstrate that fibrinogen degradation is not restricted to sites of direct plasma membrane-ECM contact, where MT1- MMP activity is concentrated, but also involves secreted proteolytic factors. Interestingly, fibrinogen degradation occurs predominantly at the macrophage rear, whereas cell-associated degradation is mainly localized at the vicinity of invadosomes. This suggests that macrophages rely on at least two spatially distinct modes of ECM degradation that may act in a complementary manner. Thus, ECM degradation by macrophages appears to involve both invadosome-associated proteolysis at the plasma membrane and extracellular proteolysis mediated by secreted factors that may remain deposited within the ECM or diffuse over short distances from the cell. This spatial organization of ECM degradation in macrophages is reminiscent of the osteoclast model, in which invadosomes form the sealing zone, spatially separated from the ruffled border where acidification and cathepsin secretion drive matrix degradation^28,29^. The relative balance between these degradation modes may determine whether ECM proteolysis primarily supports directional migration and the formation of degradation tracks or instead generates broader, more isotropic matrix remodeling. Excessive or poorly spatialized degradation could locally reduce matrix integrity and thereby alter macrophage migration or confinement. Whether such a mechanism contributes to the relatively stationary behavior observed for certain tissue-resident macrophage populations in vivo remains to be determined^30^.

Finally, the description of this long-distance degradation mode may also help to revisit some key aspects of cell invasion. Indeed, numerous works showed that nucleus passage through ECM is a key limiting factor of migration and invasion that has also consequence on cell identity^5,31–33^. Moreover, forced nuclear passage induces nuclear membrane rupture leading to DNA breakage and loss of DNA integrity. Thus, long-distance degradation could both facilitate nucleus passage by the cells able to develop this ability as macrophages but also be a mechanism to protect their DNA content from the invasion process.

### Revisiting cathepsins functions in macrophage invasion and their interplay with MMPs activity

Our quantitative analysis further supports the existence of genuine long-distance ECM degradation in macrophages. The degraded area progressively expanded beyond the cell boundary, reaching approximately 3 µm from the cell edge (for both sites) in activated OS macrophages and up to 9-10 µm (both sites) in activated OH macrophages. HCK activation also increased the normalized degradation expansion rate from approximately 0.039 h□¹ in OS to 0.063 h□¹ in OH macrophages. Interestingly, computational reaction-diffusion models based on this value predict a characteristic range of MMP-mediated ECM degradation of approximately 10 µm and an outward degradation front of ∼1 µm/min^34^. Although these parameters cannot be directly compared with our normalized expansion rate, the predicted spatial range closely matches the 9-10 µm long-distance degradation observed upon HCK activation, supporting the spatial propagation of extracellular proteolytic activity.

Interestingly, inhibition of either MMPs or cathepsins reduced fibrinogen degradation, although cathepsin inhibition produced a stronger effect, particularly in OH-activated macrophages. This suggests that both protease families contribute to fibrinogen remodeling and may act cooperatively rather than through completely independent pathways. Both classes of extracellular proteases have broad and partially overlapping substrate repertoires, including collagen, fibronectin, and fibrinogen^27,35^. Recent development of the PACMAN algorithm has further helped define protease-substrate specificities and predicted distinct cleavage kinetics among different protease families, including MMPs and cathepsins^36^. Such differences in activity, together with their distinct spatial regulation, could contribute to the localized MMP-dependent and broader cathepsin-dependent degradation patterns observed here. This functional interplay is further supported by the observation that OH activation partially preserved fibrinogen degradation in the presence of the MMP inhibitor GM6001, suggesting that enhanced cathepsin activity can partially compensate for reduced MMP activity. In addition, cathepsins may influence MMP activity through proteolytic activation of secreted MMP precursors^37^, providing a potential mechanism for crosstalk between both proteolytic systems. A major distinction between these pathways is the strong dependence of many cathepsins on acidic conditions, which can be locally regulated through the fusion of acidic vesicles or the activity of proton transporters such as the Na+/H+ exchanger NHE1.

A functional relationship between MMPs and cathepsins has also been described in osteoclasts, where combined loss of MMP9 and cathepsin K produces more pronounced defects in bone matrix remodeling than loss of either protease alone^38^. The cooperation of proteases with complementary spatial regulation and substrate preferences may therefore be particularly important for remodeling complex three-dimensional ECMs, in which fibrillar networks coexist with globular and non-fibrillar matrix components.

### HCK signaling controls an unique exo-endocytosis coupling mechanism in macrophage

OH activation revealed that exocytosis of CD63-positive vesicles can be both regulated and spatially coordinated with clathrin endocytic hotspots. Activated OH promotes CD63- positive exocytic events, as revealed by CD63-pHluorin bursts (Fig. 4), while previous work demonstrated its ability to increase endocytic events in some clathrin structures^39^. Inhibition of membrane fission of endocytosis with jasplakinolide preserved the strong association between OH and CD63-pHluorin-positive compartments but abolished their characteristic transient fusion dynamics, leading instead to their accumulation at the plasma membrane. Finally, CLEM data revealed the close spatial proximity of OH/clathrin domains with CD63- and VAMP8-positive membrane structures (Fig.5), further supporting the coexistence of endocytic and exocytic machinery within the same subcellular region. Coupling of exo- and endocytosis has been proposed for many years but remains difficult to visualize directly. Much of the evidence derives from high-temporal-resolution capacitance measurements combined with carbon-fiber amperometry in chromaffin cells, as well as studies of synaptic vesicle recycling, where exo-endocytic coupling contributes to maintaining vesicle pool composition and membrane homeostasis at presynaptic terminals^40–42^. However, the spatial organization of this coupling at defined subcellular domains remains less well characterized. Our data provide a means to visualize and experimentally perturb this coordination in macrophages, where it is associated with long-distance fibrinogen degradation. The functional significance of this physical proximity remains unclear. One possibility is that coupling the fusion of large CD63-positive endolysosomal compartments to nearby clathrin-mediated endocytosis provides a mechanism to tightly control local membrane addition and acidic cargo release while rapidly recovering excess plasma membrane. Further correlative and dynamic ultrastructural studies will be necessary to capture the transient intermediates underlying this process.

Thanks to optogenetics, it was possible to show that distinct modes of ECM degradation can coexist and be regulated in macrophages. These findings provide a framework to reinterpret pathological and clinical contexts in which both MMPs and cathepsins are present but their coordinated spatial functions have not been considered. This highlight the importance of distinguishing between invadosome-associated degradation and HCK-dependent long-distance ECM degradation when considering strategies aiming to manipulate either macrophage invasion diseases or other pathology of invasion of non-immune cells such as metastasis.

***Figure S1. Dynamic degradation of fibrinogen and gelatin by macrophages*** (A) Workflow of the quantification pipeline for degradation dynamics.

(B-C) Separation of normalized degradation fractions (%) and degradation rates (µm²/min) for all analyzed cells. Data are presented as mean ± SD. Between 12-20 cells per condition were analyzed. Statistical significance was assessed using a nonparametric Kruskal-Wallis ANOVA test.

(D) Representative intensity of degradation graphs of macrophages over fibrinogen or gelatin, separated in the different modes of degradation, showing lower intensity of degradation associated with long distance mode.

(E) Comparison of ECM degradation rates obtained using cell boundaries segmented from transmission images or membrane-labeled RAW-CAAX macrophages. Between 20 cells analyzed. Statistical significance was assessed using an unpaired Welch’s t-test.

(F) Time-lapse imaging of the classification. The white line indicates invadosome overlay associated with degradation on fibrinogen and gelatin.

(G) Temporal progression of degradation at the same sites is shown (normalized mean intensity as a function of time). Invadosome formation preceded the progressive loss of fibrinogen fluorescence, illustrating the temporal relationship between invadosome activity and local ECM degradation.

(H) Time-lapse imaging of the classification associated with either the cell or the invadosome.Frames were acquired every 4 minutes for up to 16 hours. Scale bar: 20 µm.

***Figure S2. Comparative degradation dynamics of MDA cancer cells and macrophages on fibrinogen and gelatin substrates*** (A) Time-lapse imaging of MDA cells on fibrinogen or gelatin. The white line indicates lamellipodia or invadopodia overlay associated with degradation. Frames were acquired every 10 minutes for up to 16 hours. Scale bar: 20 µm.

***Figure S3. Role of Src family kinases in macrophage degradation dynamics*** (A-B) Separation of normalized degradation fractions (%) and degradation rates (µm²/min; mean ± SD.) for all analyzed cells. Data are presented as mean ± SD. Between 20-30 cells per condition were analyzed. Statistical significance was assessed using a nonparametric Kruskal-Wallis ANOVA test.

(C) Representative intensity of degradation graphs of macrophages expressing OS or OH over fibrinogen, separated in the different modes of degradation.

(D-E) Representative time course of the minor axis (width) and major axis (length) of the cell and corresponding degraded area in OS- and OH-expressing macrophages.

***Figure S4. Secretome profiling of OS- and OH-activated macrophages reveals distinct ECM-remodeling and lysosomal/vesicular signatures***. (A) Secretome analysis of proteins differentially secreted in stimulated WT versus OS and WT versus OH macrophages (p ≤ 0.05, log□FC ≥ 1). Data was obtained from triplicates after 24 h stimulation in serum-free conditions.

(B) Venn diagram of enriched proteins shared and uniquely detected in OS and OH.

(C) GO enrichment analysis of cellular components (g:Profiler) associated with secretion.

***Figure S5. Parameters and controls of the PhysiCell computational model of ECM degradation.*** (A) Spatial positioning of proteolytic components in the migrating-cell model. MMPs and cathepsins were released in association with the cell, whereas H+ release was spatially biased toward the cell rear according to the migration direction.

(B) Parameters used for substrate diffusion, secretion, decay, cell–ECM interactions, and cathepsin/H+-dependent ECM degradation in the PhysiCell model. Basal diffusion coefficients for H+ and cathepsins were set to 10 µm²/min, with secretion rates of 0.5 unit/min for both components.

(C) Representative 24-h control simulation of a non-migrating, non-degrading cell, showing the absence of spontaneous ECM degradation in the model.

## Material and methods

### Experimental cell models and culture

Murine RAW 264.7 macrophages cell line was transiently and/or stably transduced using a viral and/or transposon strategy. Cells were cultivated at 37□C and 5% CO2 in RPMI 1640 medium supplemented with GlutaMAX™ (Gibco, Thermo Fisher Scientific), 10% fetal bovine serum (FBS) and 1% (v/v) penicillin-streptomycin (P/S). Transient transfection of RAW 264.7 macrophages was performed using jetPRIME® transfection reagent (Polyplus-transfection) with optimized conditions (0.8□ µg plasmid DNA, 75□µL jetPRIME® buffer, and 1.6□µL reagent, in a final volume of 700□µL).

Epithelial human breast cancer MDA-MB 231 cells were cultivated at same conditions in DMEM, high glucose, GlutaMAX™ supplement (Gibco, Thermo Fisher Scientific) with 10% FBS and 1% P/S.

### DNA constructs and plasmids

In this study, the various Optogenetic SFKs plasmids are the result of C-terminal fusion of CRY2-mCherry to the indicated SFK mutants (Torres-Torres, Rivier et al., 2025). These, and additional recombinant DNA constructs are listed in key resource table.

Plasmids were constructed using Gibson Assembly, according to the manufacturer’s instructions (New England Biolabs, NEB), made nicely available by the indicated principal investigator through purchased from Addgene (see details articles and addgene number in key resource table) or kindly given as gifts by the indicated collaborators.

### Obtention of stabling expressing RAW 264.7 macrophages

#### Lentiviral infection

Lentiviruses were produced by co-transfecting pC57GPBEB GagPol MLV, pSUSVSVG (gifts of Dr Negre, ANIRA Platform, SFR Biosciences, Lyon, France) and each plasmid of interest using Lipofectamine2000 (Invitrogen) in HEK293 FT cells (gift of Dr Negre) plated in 6-well plates at 50% confluency. The medium was changed 24 h later. The viral supernatant was collected after 72 h and was filtered with 0.45 μm filters. RAW 264.7 cells were plated in 6-well plates so as to achieve 60% confluency on the day of infection. The filtered supernatant was directly used to infect cells of interest. The medium was changed 24 h after infection. After 10 days of decontamination, cells were FACS sorted (Aria cell sorter 2000, BD) based on the level of expression of mCherry-tagged OSFK in order to have no dark binding of the CRY2 system, using excitation with a 561 nm LASER.

#### Transposon

GFP-tagged endocytic markers were stably expressed in RAW 264.7 cell lines stably expressing optoSFKs using the Sleeping Beauty transposon system. Briefly, cells were seeded one day prior to transfection, at approximately 50% confluency. Transfection was performed following the previously mentioned jetPRIME protocol, in antibiotic-free RPMI medium. Plasmid DNA encoding the marker of interest was co-transfected with the Sleeping Beauty transposase plasmid at a 1:10 ratio (total DNA). After 5-6 hours, the transfection medium was replaced with complete RPMI medium supplemented with 10% FBS and 1% P/S. Puromycin selection (5□µg/mL) was initiated 48 hours post-transfection and maintained by replacing the medium every 2 days until all non-resistant cells were eliminated.

### Degradation matrix and imaging

#### Fixed degradation assays

Sterile 12-mm coverslips were placed into 24-well plates, sterilized with 70% ethanol, and plasma cleaned. Coverslips were rinsed once with sterile H□O, coated with 500 µL poly-L-lysine (50 µg/mL; Sigma-Aldrich, P4832) for 20 min at room temperature, rinsed twice with H□O and once with PBS, and then crosslinked with glutaraldehyde (0.5% v/v in PBS; Sigma-Aldrich, cat. F1056) for 15 min on ice. After three washes with cold PBS, a fibrinogen mixture consisting of non-fluorescent fibrinogen (0.2 mg/mL) and fluorescent fibrinogen (0.5 µg/mL; Thermo Fisher Scientific, Cat. No. F13191) was applied. Same protocol was used for gelatin^4^. For coating, 40 µL of the fibrinogen mix was placed on parafilm, and coverslips (poly-L-lysine/glutaraldehyde side down) were incubated on the droplet for 10 min at room temperature. Coverslips were then rinsed three times with PBS and returned to 24-well plates. RAW 264.7 macrophages were seeded and cultured in complete RPMI medium (10% FBS, 1% P/S). After incubation, cells were fixed with 4% or 8% paraformaldehyde (PFA) and processed for imaging.

#### Live degradation assays

For live-cell imaging, coatings were prepared directly on glass-bottom chambers (Ibidi 4-, 8-, or 24-well plates). A custom 3D-printed stamp was used to ensure homogeneous distribution of the matrix over the glass surface. Chambers were coated with the fibrinogen mixture as described above. After seeding, cells were imaged under live conditions using time-lapse microscopy to monitor dynamic matrix degradation.

#### Confocal imaging

Live and fixed imaging were performed using a Confobright inverted confocal microscope (Nikon Eclipse Ti2, scanner A1R; Nikon Instruments) equipped with the NIS-Elements software and adaptive optics (ALPAO) for aberration correction. The system was fitted with a fast resonant scanner for high-speed time-lapse imaging and an Okolab incubation chamber maintained at 37 °C for live-cell experiments. Fluorescence excitation was achieved using 405, 488, 561, and 640 nm lasers, with corresponding emission filters for DAPI, FITC, Rhodamine, and Cy5 channels. The following Nikon objectives were used depending on the experiment: Plan Apo 20×/0.75 for fixed conditions, and Plan Fluor 40×/1.30 Oil (WD 0.24 mm) zoom 2x for live conditions. For long-term live imaging, the Perfect Focus System (PFS) was used to maintain focus stability over extended acquisitions (16-24 h) minimal drift.

### Single cells optogenetics experiments and TIRF imaging

Live-cell imaging was performed on an inverted motorized microscope (Zeiss AxioVert 200M) equipped with a Coolsnap HQ2 camera (Photometrics), 100× (NA 1.46, oil) Plan-Apochromat oil objective (Rapp Optoelectronic). TIRF illumination was performed using DPSS lasers at 488 nm, and 561 nm, combined with a Slider TIRF2 module (ZEISS). During imaging, cells were placed on a heated 37°C stage (ZEISS) combined with an incubator of CO2 (XL incubator, PeCon). Acquisition and experimental control were managed with MetaMorph software (Universal Imaging).

Image brightness and contrast were adjusted post-acquisition using ImageJ to enhance visualization of optogenetic probe recruitment. For photo-stimulation experiments, brightness settings were standardized across conditions and optimized based on the maximal intensity observed in the stimulated images. This adjustment was applied uniformly to allow accurate comparison and to highlight probe dynamics upon activation.

### Secretome

RAW 264.7 macrophages (2 × 10□cells/well) were seeded in 6-well plates and allowed to adhere overnight in complete RPMI medium (10% FBS, 1% P/S). The following day, cells were washed four times with PBS to remove serum and incubated in 1.5 mL serum-free RPMI for 24 h, with optogenetic stimulations performed every 4 min for opto-expressing cells. Supernatants were collected and clarified by centrifugation at 1,200 rpm for 10 min at 4 °C to remove cells, followed by centrifugation at 4,000 rpm for 10 min to eliminate debris. Secreted proteins were concentrated using Amicon Ultra-2 centrifugal filters (3 kDa cutoff; Merck Millipore, UFC200324) preconditioned with 500 µL of 1 mg/mL BSA in sterile ddH□O, centrifuged at 4,000 rpm for 20 min at 4 °C, and rinsed three times with 1 mL ddH□O. Clarified supernatants were loaded onto the filters and centrifuged at 4,000 rpm for 40 min at 4 °C. Concentrates were recovered by inverting the filters and centrifuging at 1,000 rpm for 1-2 min. Samples were snap-frozen and kept at -80 °C until analysis. Columns were rinsed with ddH□O and stored in ddH□O containing 0.1% NaN□at 4 °C.

Mass spectrometry analysis identified approximately 2,400 proteins, and statistical processing was performed using ProStaR software. Protein abundances were log□ - transformed, normalized, and missing values imputed. Proteins were considered differentially secreted when showing a log□(fold change) ≥ 1 or ≤ −1 (fold change > 2) and a p ≤ 0.05 (limma test).

### PREM and CLEM Electron Microscopy

Adherent plasma membranes of murine RAW 264.7 macrophage cell lines on glass coverslips were disrupted by sonication according to Heuser (2000b). The unroofed cells were then fixed and processed as previously described (Vassilopoulos et al. 2019). Sequential treatments involved 0.55% OsO4, 1% tannic acid, and 1% uranyl acetate, followed by graded ethanol dehydration and substitution with hexamethyldisilane (Sigma). Once dried, the samples were rotary shadowed with 2 nm of platinum and 6 nm of carbon using a high-vacuum sputter coater (Leica Microsystems). The resulting platinum replica was removed from the glass with 5% hydrofluoric acid, thoroughly washed in distilled water, and mounted on 200 mesh formvar/carbon-coated EM grids. The images were captured with a digital camera (Xarosa) attached to a transmission electron microscope (Joel, USA) operating at 120 kV, with the grids mounted on an eccentric side-entry goniometer stage. The images were adjusted for brightness and contrast using Adobe Photoshop (Adobe, USA) and presented in inverted contrast. For CLEM, unroofed and stained samples were imaged by spinning disk microscope, prior to make the replica. Grids were first imaged by low-magnification EM to relocate the light microscopy-imaged area, followed by high-magnification EM of the corresponding ROIs, and aligned using affine transformation via the eC-CLEM plugin in ICY(Paul-Gilloteaux, et al., 2017).

### Computational Framework for simulation of MMP, cathepsins and acid interplays for simulation of migration/ECM degradation in silico

All simulations were performed using PhysiCell (version 1.14.0)^43,44^, an open-source, agent- based modeling framework for multicellular systems. To investigate extracellular matrix (ECM) degradation, we implemented a custom model introducing two alternative degradation mechanisms: (1) ECM breakdown mediated by membrane-bound matrix metalloproteinases (MMPs), and (2) degradation driven by Cathepsin secretion and acidification via hydrogen ions (H⁺). In the first case, cells degrade ECM directly at the leading edge through localized MMP activity. In the second mechanism, Cathepsin is secreted at the cell front while H⁺ ions are released at the rear, creating an acidic microenvironment that activates Cathepsin and promotes ECM proteolysis.

All custom functionalities were implemented in the custom.cpp file without modifying PhysiCell’s core source code. Two major extensions were introduced: (i) cells can secrete or uptake substrates not only within their own voxel but also in voxels located in front of and behind the cell along its migration vector, allowing for spatially resolved degradation; and (ii) a system of ordinary differential equations (ODEs) governing ECM degradation dynamics was added. This system accounts for the interaction between ECM, Cathepsin, and H⁺ concentrations, enabling simulation of proteolytic and acidification-driven degradation processes over time and space.

#### Model Setup and Parameters

Simulations were performed in a 2D environment containing a single motile cell surrounded by ECM. The cell moved with a random walk with a persistence time of 60 minutes and a base velocity of 1 µm/min. Cellular proliferation, growth, and death processes were deactivated to focus exclusively on motility and matrix degradation dynamics.

The model included diffusion and decay of ECM, H⁺, and Cathepsin, as well as reaction terms for their biochemical interactions. Reaction rates followed the law of mass action for a third-order reaction:

r=kreaction[ECM][H+][Cathepsin]

where kreaction is the rate constant (set to 100 in dimensionless units). Depending on simulation parameters, H⁺ and Cathepsin could be optionally consumed during the reaction. Concentration updates were computed using explicit Euler integration with a time step of 0.01 min.

Key biophysical parameters were chosen to match experimental scales or default PhysiCell values: diffusion coefficients for H⁺ and Cathepsin were both set to 10 µm²/min, secretion rates to 0.5 unit/min, and ECM uptake rate to 0.2 unit/min. Adhesion and repulsion between the cell and ECM were parameterized by dimensionless strengths of 1.0 and 5.0, respectively. Parameter toggles controlled the activation of degradation modes (MMP14, cathepsin, consume_H, and consume_cathepsin).

#### Implementation and Reproducibility

The full source code and configuration files are publicly available at https://github.com/marcorusc/pc4_degradation.

The model can be compiled within PhysiCell by placing the degradation project folder in Sample_projects, replacing the default Makefile, and running make degradation; make -j. The resulting executable is compatible with PhysiCell-Studio for visualization and parameter tuning. All simulations were initialized with uniform ECM distribution and periodic random migration of the cell. Different parameter combinations were explored to assess the relative contributions of MMP- and Cathepsin-mediated degradation and to study the spatial and temporal effects of acidification, diffusion, and substrate consumption on ECM remodeling.

**Table 1.**
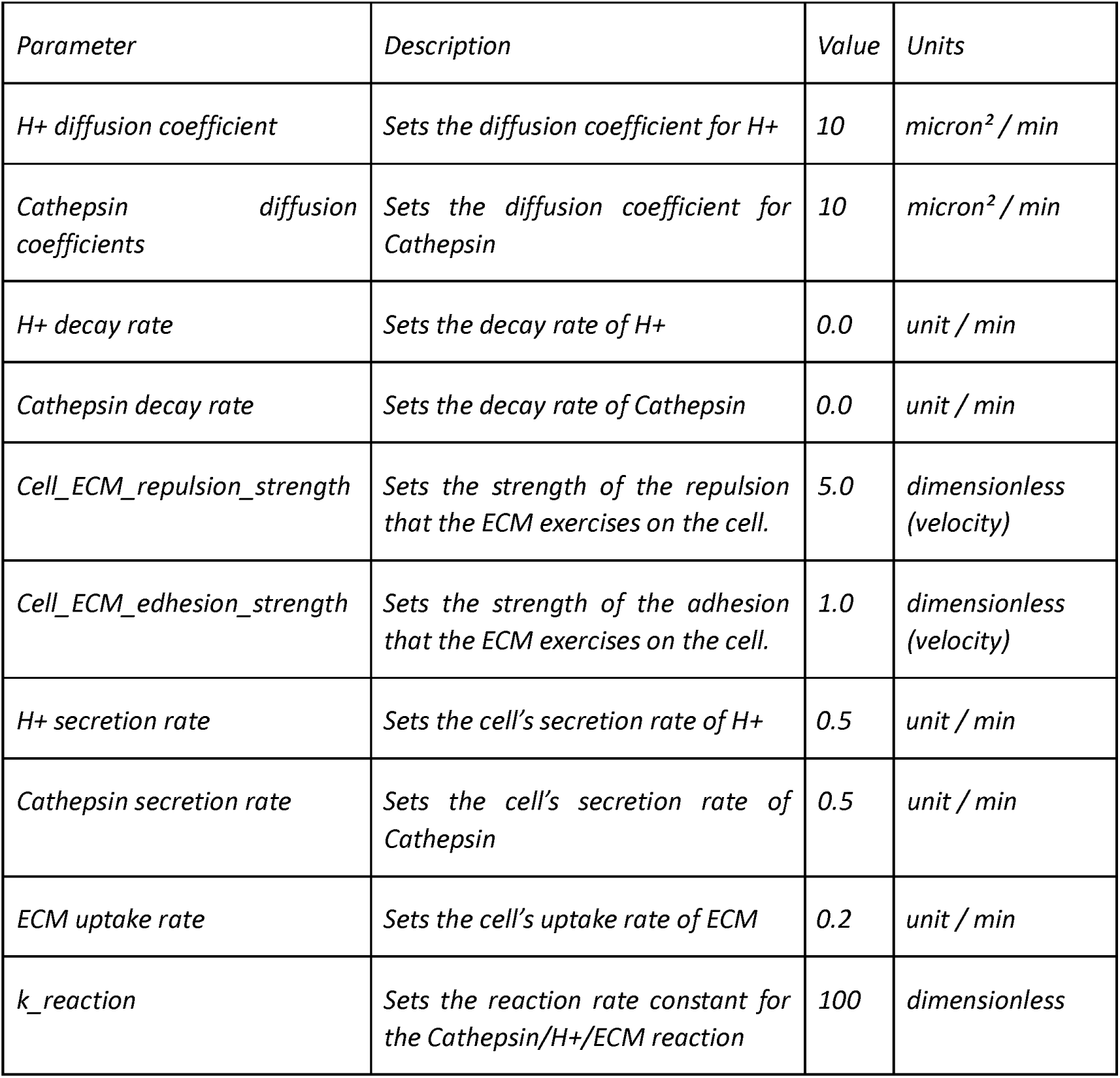
List of parameters introduced in the model to control the secretion, uptake, and decay of the substrates, including the adhesion and repulsion strength of the ECM and the reaction parameter for the ODE system.

**Table 2.**
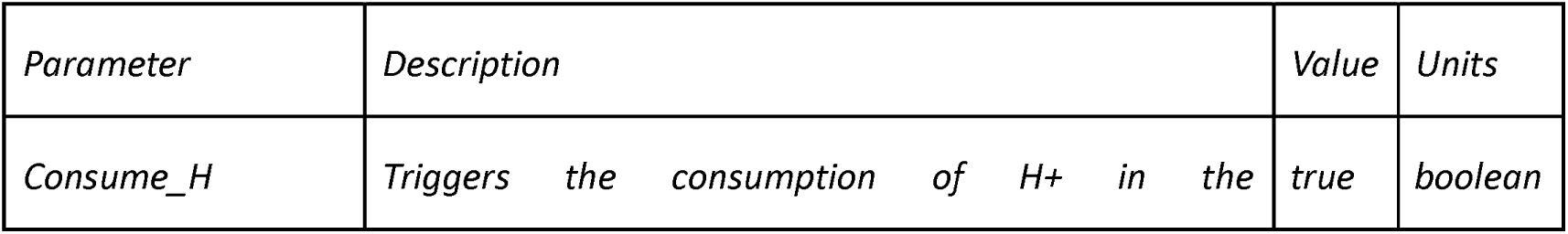
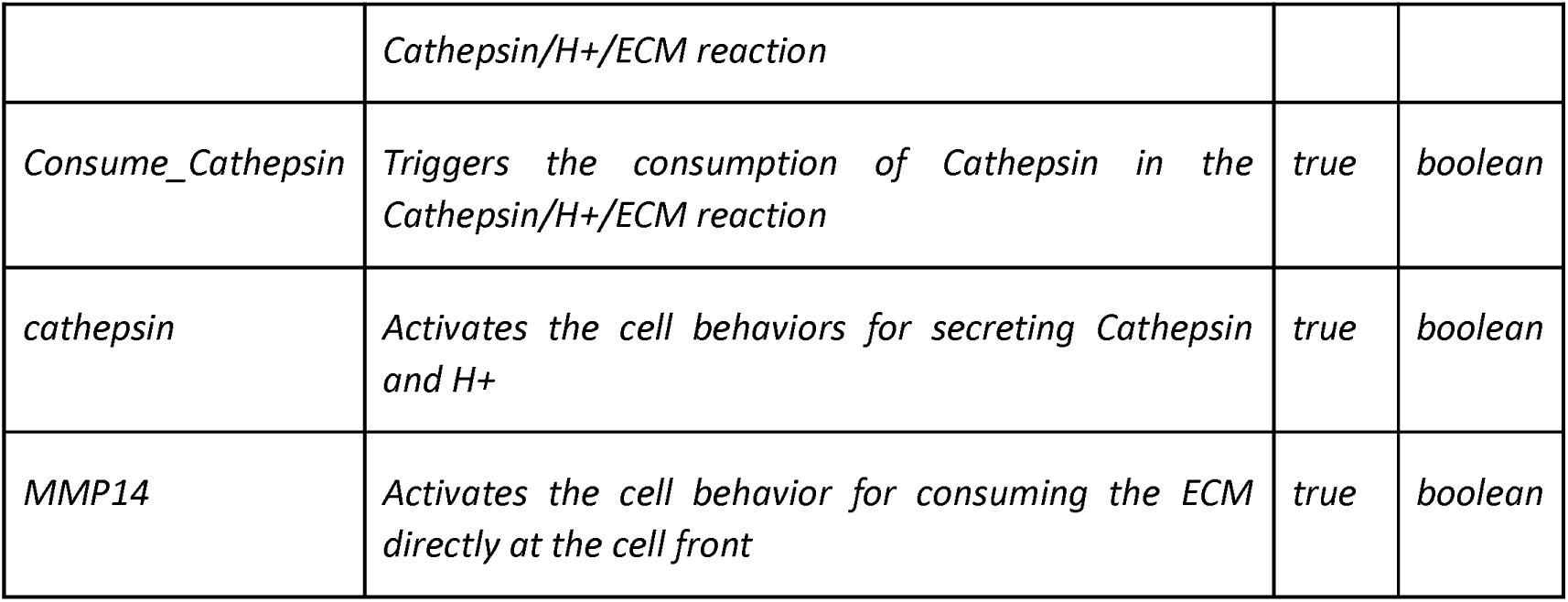
List of parameters that control the switch between the degradation modes.

### Image analysis

#### Co-occurrence analysis

Co-occurrence analysis was performed using the ComDet plugin (Eugene Katrukha / https://github.com/UU-cellbiology/ComDet/wiki / in ImageJ. The plugin was configured to detect spots based on a defined particle size and intensity threshold. Colocalization events were identified based on spatial proximity between detected puncta in the two channels within a given maximal distance (3-4 pixels). The percentage of co-occurring spots was calculated relative to the total number of spots in each channel.

#### Live-degradation analysis

Segmentations of degradation areas were generated using Ilastik machine learning software^45^. Segmented images were subsequently quantified with a custom Fiji macro. Matrix degradation was expressed as the percentage of degraded area relative to the total imaged surface.

#### Degradation expansion analysis

For the geometrical analysis of cell and degradation areas, segmented binary images were analyzed in Fiji using the Analyze Particles function with the Fit Ellipse option. The resulting major and minor axis lengths, together with ellipse orientation, were extracted from the Results table. The minor axis length was plotted as a function of time to quantify the temporal expansion of the degraded area. A linear regression was fitted to each trajectory, and the slope of the regression was used as the degradation expansion rate. For visualization, the fitted ellipse parameters were used to reconstruct and display the major and minor axes of the cell and degraded areas.

### Statistical analysis

Statistical analyses were performed using GraphPad Prism 10 (GraphPad Software). Statistical significance was defined as P < 0.05. Specific statistical tests used are indicated in the figure legends. Data are presented as mean ± standard deviation (SD), as specified. Significance levels are denoted as follows: ns, not significant (P□>□0.05); *P < 0.05; **P□<□0.01; ***P□<□0.001; ****P□<□0.0001.

Schematic figures were created using BioRender. Statistical analyses and graph generation were performed using GraphPad Prism (version 10). Image analysis was done in Fiji^46^ with segmentation performed in Ilastik^45^. Final figure assembly and image formatting were carried out in Adobe Illustrator. The main text was written by CTT and OD, then submitted to Mistral AI and ChatGPT for wording optimization.

## Acknowledgements

This work was funded by ANR “NODES” and ARC programs (coordinator: O.D.), by LLNC for PhD funding of C.T.T., P.R. and L.C. and by AC. 4^th^ of PhD of C.T.-T. was funded by ARC. We thank the strong and efficient support of the imaging platform MicroCell being part of FBI AURA node. We acknowledge France-BioImaging infrastructure supported by the French National Research Agency (ANR-24-INBS-0005 FBI BIOGEN).

## Authors Contributions

C.T.-T., L.C., C.O. and O.D. generated constructs and cell lines, performed and analyzed data in cell biology. C.T.-T., L. C.-P., A.P. B. and F.S. developed co-occurrence, ECM degradation quantification and elliptic analysis. B.G. and L.H. developed spatially controlled activation in segmented region of interests (IN or OUT podosomes). A.G. support the excellence of TIRF imaging on the MicroCell platform. C.T.-T., and O.D designed the study, and C.T.-T., and O.D. wrote the paper with significant contributions from all authors.

## Notes

### Competing Interest Statement

The authors have declared no competing interest.

