## Supplementary material for "Hck signaling drives long-distance ECM degradation through endo-exocytosis coupling in macrophages": Sup Fig 1 to 5

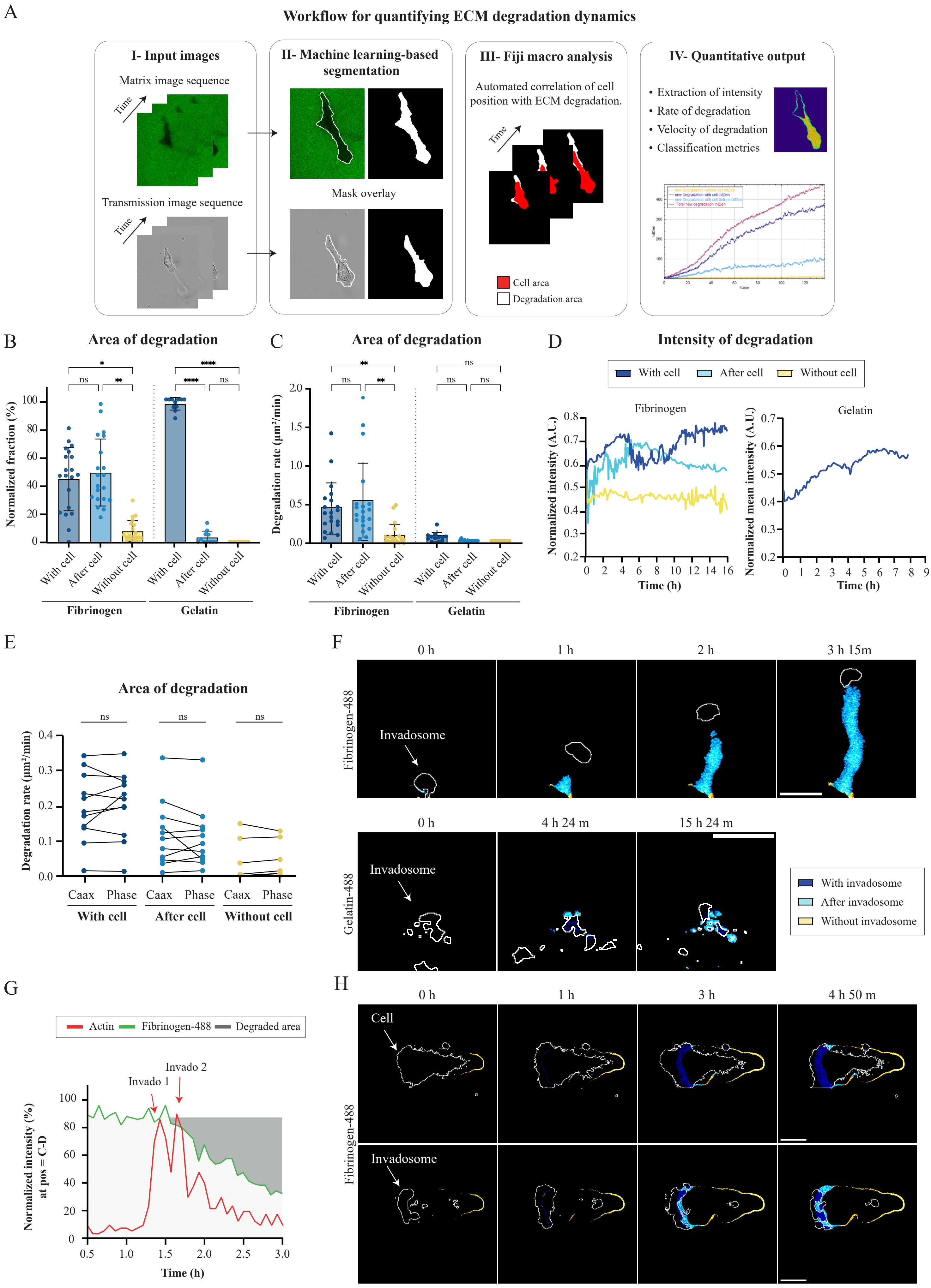
A
Workflow for quantifying ECM degradation dynamics
I- Input images
Matrix image sequence
Time
Transmission image sequence
II- Machine learning-based segmentation
Mask overlay
III- Fiji macro analysis
Automated correlation of cell position with ECM degradation.
Cell area
Degradation area
IV- Quantitative output
• Extraction of intensity
• Rate of degradation
• Velocity of degradation
• Classification metrics
B
Area of degradation
Normalized fraction (%)
With cell
After cell
Without cell
Fibrinogen
Gelatin
C
Area of degradation
Degradation rate (µm²/min)
D
Intensity of degradation
With cell
After cell
Without cell
Fibrinogen
Gelatin
Normalized intensity (A.U.)
Normalized mean intensity (A.U.)
Time (h)
E
Area of degradation
Degradation rate (µm²/min)
Caax
Phase
With cell
After cell
Without cell
F
0 h
1 h
2 h
3 h 15m
Fibrinogen-488
Invadosome
0 h
4 h 24 m
15 h 24 m
Gelatin-488
With invadosome
After invadosome
Without invadosome
G
Actin
Fibrinogen-488
Degraded area
Invado 1
Invado 2
Normalized intensity (%) at pos = C-D
Time (h)
H
0 h
1 h
3 h
4 h 50 m
Cell
Invadosome
Fibrinogen-488

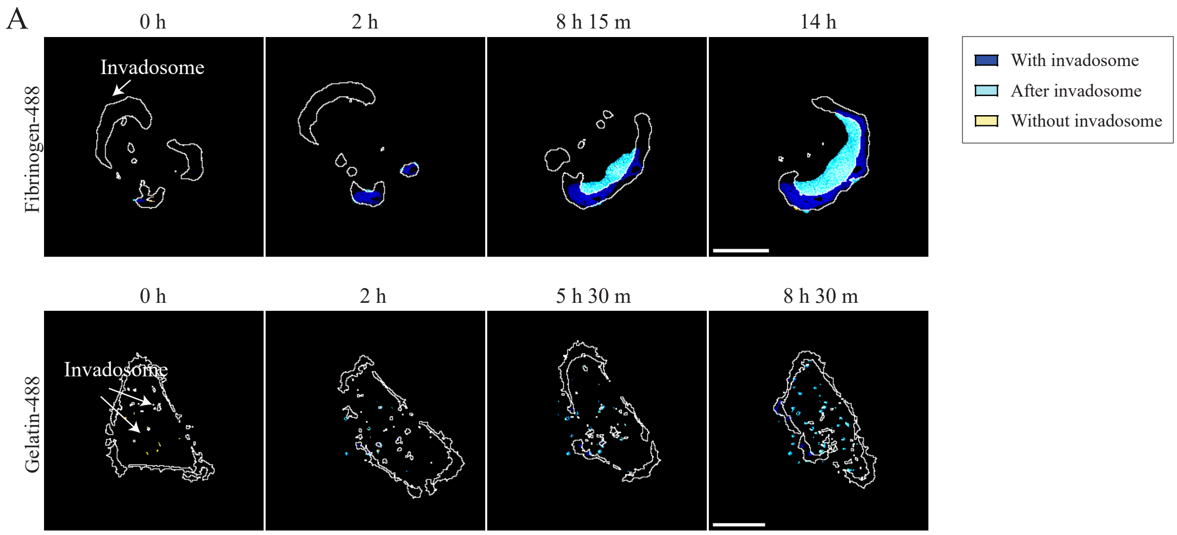
A
0 h
2 h
8 h 15 m
14 h
Fibrinogen-488
Invadosome
0 h
2 h
5 h 30 m
8 h 30 m
Gelatin-488
Invadosome
With invadosome
After invadosome
Without invadosome

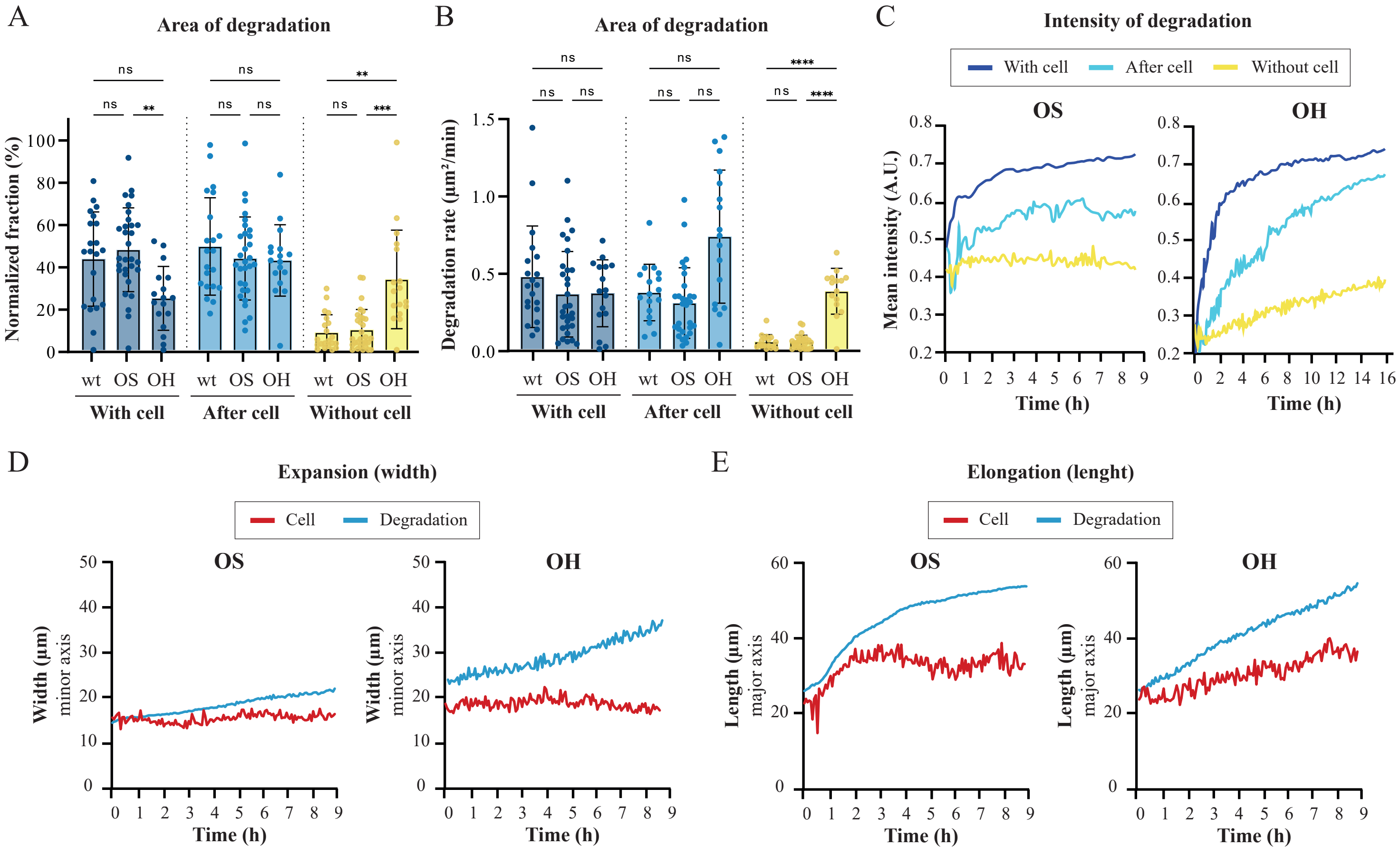
A
Area of degradation
Normalized fraction (%)
ns
**
***
wt OS OH
With cell
After cell
Without cell
B
Area of degradation
Degradation rate (µm²/min)
****
C
Intensity of degradation
With cell
After cell
Without cell
OS
OH
Mean intensity (A.U.)
Time (h)
D
Expansion (width)
Cell
Degradation
Width (µm)
minor axis
E
Elongation (lenght)
Length (µm)
major axis

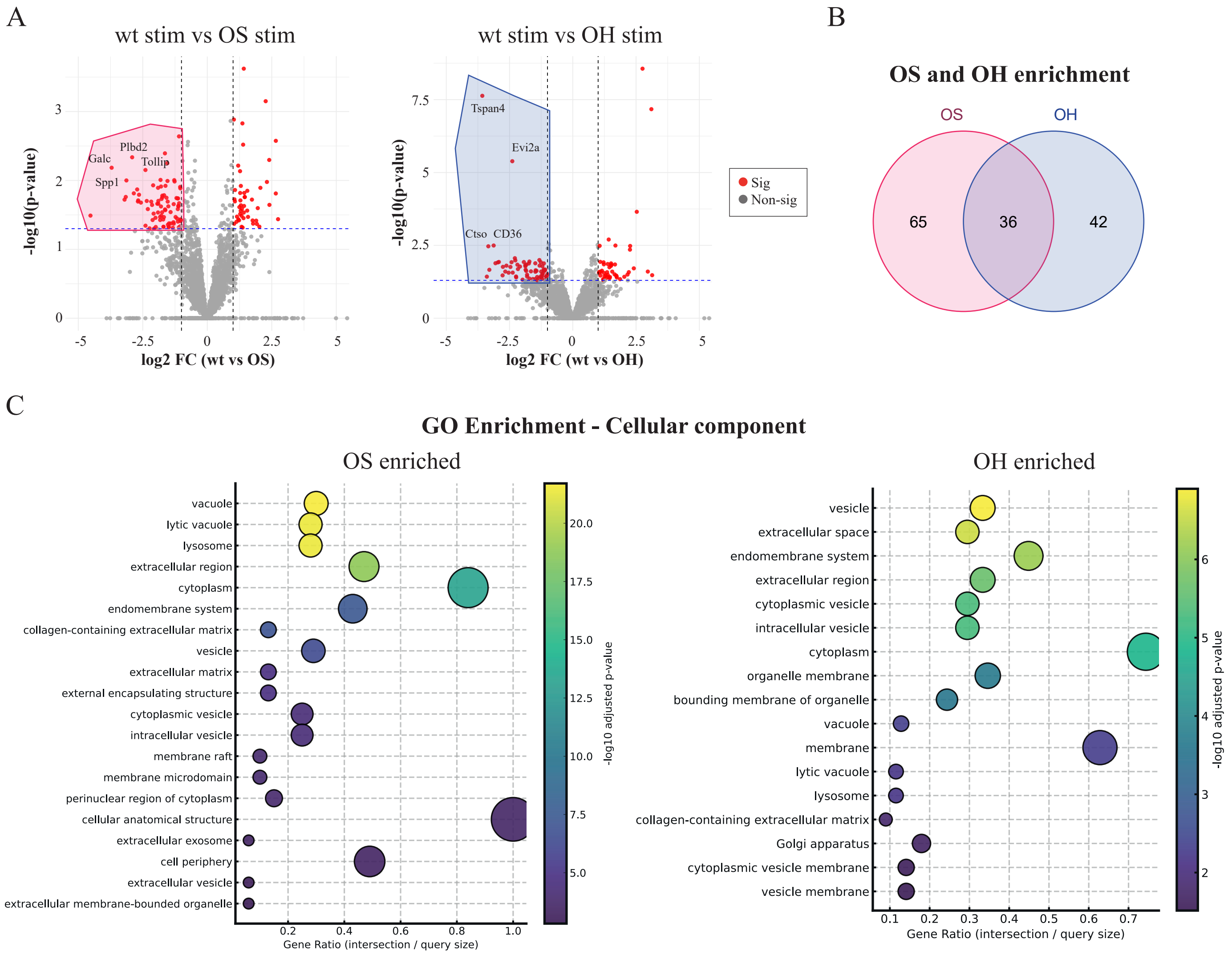
A
wt stim vs OS stim
-log10(p-value)
log2 FC (wt vs OS)
Plbd2
Galc
Tollip
Spp1
wt stim vs OH stim
-log10(p-value)
log2 FC (wt vs OH)
Tspan4
Evi2a
Ctso
CD36
Sig
Non-sig
B
OS and OH enrichment
OS
OH
65
36
42
C
GO Enrichment - Cellular component
OS enriched
vacuole
lytic vacuole
lysosome
extracellular region
cytoplasm
endomembrane system
collagen-containing extracellular matrix
vesicle
extracellular matrix
external encapsulating structure
cytoplasmic vesicle
intracellular vesicle
membrane raft
membrane microdomain
perinuclear region of cytoplasm
cellular anatomical structure
extracellular exosome
cell periphery
extracellular vesicle
extracellular membrane-bounded organelle
Gene Ratio (intersection / query size)
-log10 adjusted p-value
OH enriched
vesicle
extracellular space
endomembrane system
extracellular region
cytoplasmic vesicle
intracellular vesicle
cytoplasm
organelle membrane
bounding membrane of organelle
vacuole
membrane
lytic vacuole
lysosome
collagen-containing extracellular matrix
Golgi apparatus
cytoplasmic vesicle membrane
vesicle membrane
Gene Ratio (intersection / query size)
-log10 adjusted p-value

A

**Positioning of MMP, cathepsins and H+ on a migrative cell:**

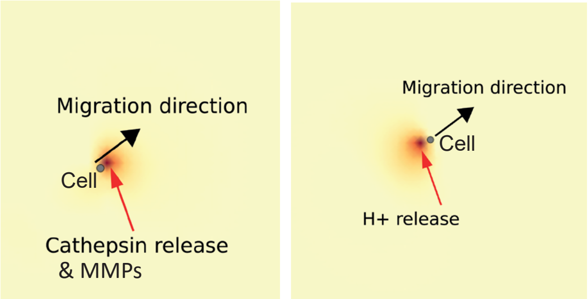


B

**Parameters of substrate rates:**

| Parameter | Description | Value | Units |
|---|---|---|---|
| H+ diffusion coefficients | Sets the diffusion coefficient for H+ | 10 | micron² / min |
| Cathepsin diffusion coefficients | Sets the diffusion coefficient for Cathepsin | 10 | micron² / min |
| H+ decay rate | Sets the decay rate of H+ | 0.0 | unit / min |
| Cathepsin decay rate | Sets the decay rate of Cathepsin | 0.0 | unit / min |
| Cell_ECM_repulsion | Sets the strength of the | 5.0 | dimensionless |
| _strength | repulsion that the ECM exercises on the cell. | | (velocity) |
| Cell_ECM_edhesion _strength | Sets the strength of the adhesion that the ECM exercises on the cell. | 1.0 | dimensionless (velocity) |
| H+ secretion rate | Sets the cell's secretion rate of H+ | 0.5 | unit / min |
| Cathepsin secretion rate | Sets the cell's secretion rate of Cathepsin | 0.5 | unit / min |
| ECM uptake rate | Sets the cell's uptake rate of ECM | 0.2 | unit / min |
| k_reaction | Sets the reaction rate constant for the Cathepsin/H+/ECM reaction | 100 | dimensionless |

C

24-h simulation of cell not degrading; not migrating

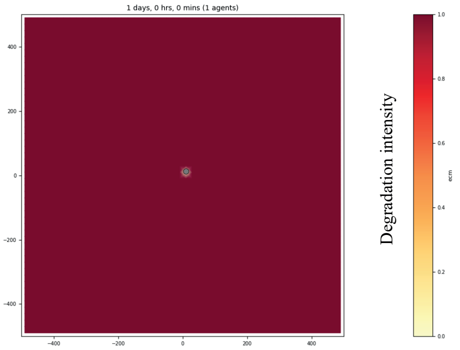
